# Structural basis of *Mycobacterium* Fluoroquinolone Resistance Protein D (MfpD), a versatile pathogeny protein from the *mfp* conservon of *Mycobacterium tuberculosis*

**DOI:** 10.64898/2026.03.03.709265

**Authors:** Antoine Gedeon, Maureen Micaletto, Daniela Megrian, Elodie Carmen Leroy, Elorri Barbier, Bertrand Raynal, Ahmed Haouz, Pedro M. Alzari, Claudine Mayer, Stéphanie Petrella

## Abstract

The *mfp* conservon of *Mycobacterium tuberculosis* has been associated with fluoroquinolone resistance and encodes five conserved proteins, including the small GTPase MfpB and its regulatory partner MfpD. In this study, we combined phylogenetic, structural, and biophysical approaches to define the molecular basis of MfpD function. MfpD adopts a Roadblock/LC7-like α/β fold and forms a stable dimer in solution, with hydrophobic α2-helix interactions stabilizing the interface. Additional biophysical analyses and AlphaFold3 modeling suggest that MfpD may promote GTP hydrolysis by MfpB through a noncanonical Switch I-dependent mechanism. These findings establish the first structural framework for MfpD-MfpB interactions, building on previously identified *in vitro* catalytic properties and proposing new insights into MfpD’s non-catalytic pathogenesis activity of MfpD in macrophages.

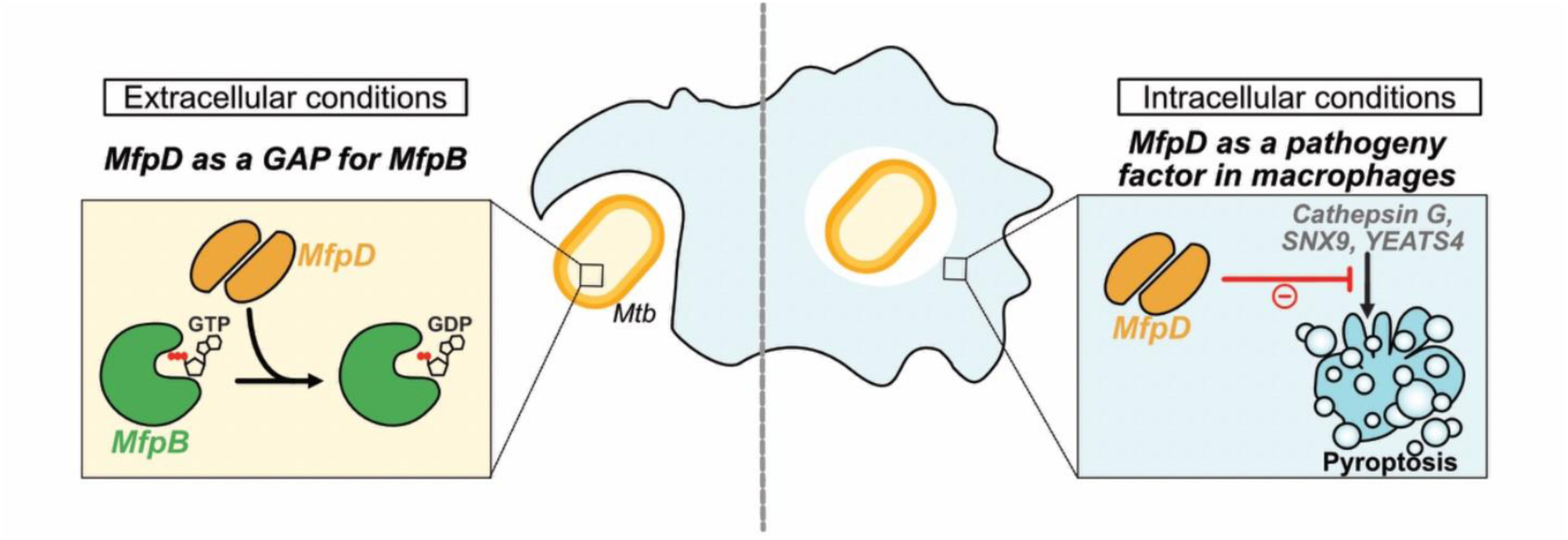
GRAPHICAL ABSTRACT.

## Introduction

Tuberculosis, a communicable airborne disease caused by *Mycobacterium tuberculosis* (*Mtb*), remains a serious threat of public health. The rapid emergence of multidrug-resistance (MDR) often leaves patients without any further treatment options^1^. One of the second regimen drug used to treat MDR strains, named fluoroquinolones (FQs), acts as a specific bacterial type II topoisomerases poison (i.e. DNA gyrase and topoisomerase IV (Topo IV))^2,3^. DNA gyrase and Topo IV are homologous heterotetrameric enzymes that share similar structural organizations but differ in activity, with one being more specialized in controlling superhelical torsions in DNA and the other being responsible for decatenation. Both enzymes are widely conserved among bacteria, except in some species such as *Mtb* where DNA gyrase is the sole type II topoisomerase, rendering it an even more attractive drug target^4,5^. From a molecular mechanistic perspective, FQs bind to two symmetric sites within the enzyme-DNA complex and block it with a covalent phospho-tyrosyl adduct between the catalytic tyrosines and the double-stranded DNA. This irreversible inhibition leads to the accumulation of double-stranded broken DNA fragments, thereby compromising genome integrity. Consequently, targeting DNA gyrase with FQs effectively disrupts bacterial DNA replication and transcriptional processes, ultimately leading to bacterial cell death^6,7^.

Nonetheless, resistance to these second line drugs has also been reported. Although this is mostly associated with the presence of mutations at the FQs binding site of *Mtb* DNA gyrase at the FQs binding sites^8,9^, overexpression in *M. smegmatis* (*Msm*) of a protein named MfpA (for “***M***ycobacterium ***F***Q resistance protein ***A***”) conferred low-level resistance to two fluoroquinolones, ciprofloxacin and sparfloxacin^10,11^. *Mtb* contains a similar 183–amino acid MfpA homolog (Rv3361c) that is 67% identical to the 192-residue *Msm* MfpA. The gene *rv3361c* (encoding MfpA in *Mtb*) is located in a conservon, termed “*mfp* conservon”, with four other coding sequences *rv3362c*, *rv3363c*, *rv3364c* and *rv3365c* as part of the same transcriptional unit^12^ (*Figure 1a*). The four corresponding proteins, named MfpB, MfpC, MfpD and MfpE, respectively, are highly conserved between *Mtb* and *Msm* (*Supplementary Table 1*). Interestingly, previous reports have revealed the existence of one to thirteen copies of *mfp* conservon-like homologs of such conservon in many Actinomycetes^13^, and It was suggested that the proteins might form a membrane-associated heterocomplex similar to eukaryotic G protein-coupled regulatory systems^13,14^. In the case of the *mfp* conservon, which is the only one identified to date in *Mtb*, the megacomplex is directly linked to DNA gyrase regulation^15^ (*Figure 1b, c*). In this scheme, MfpE, annotated as a putative membrane protein with a phosphorelay sensor histidine kinase activity, could be part of a signal transduction system that is activated in response to unknown extracellular cues that might regulate the MfpB-MfpC-MfpD molecular switch cycle. In *Msm*, MfpB acts as a GTPase and is crucial for regulating MfpA when MfpB is bound to GTP^16^. Recent findings suggest that MfpC and MfpD regulate MfpB, with MfpC acting as a guanine nucleotide exchange factor (GEF) to release GDP, and MfpD functioning as a guanine activating protein (GAP) to facilitate GTP hydrolysis by MfpB^17,18^. MfpA, identified as a pentapeptide repeat protein and a structural DNA mimic, was found to modulate DNA gyrase activity and protect it from FQs^19^. Cross-talk between MfpA and MfpB exists, as MfpB is essential for MfpA-mediated FQs resistance in living organisms and can modulate the interaction of MfpA’s interaction with DNA gyrase^8,16^.

**Figure 1.**
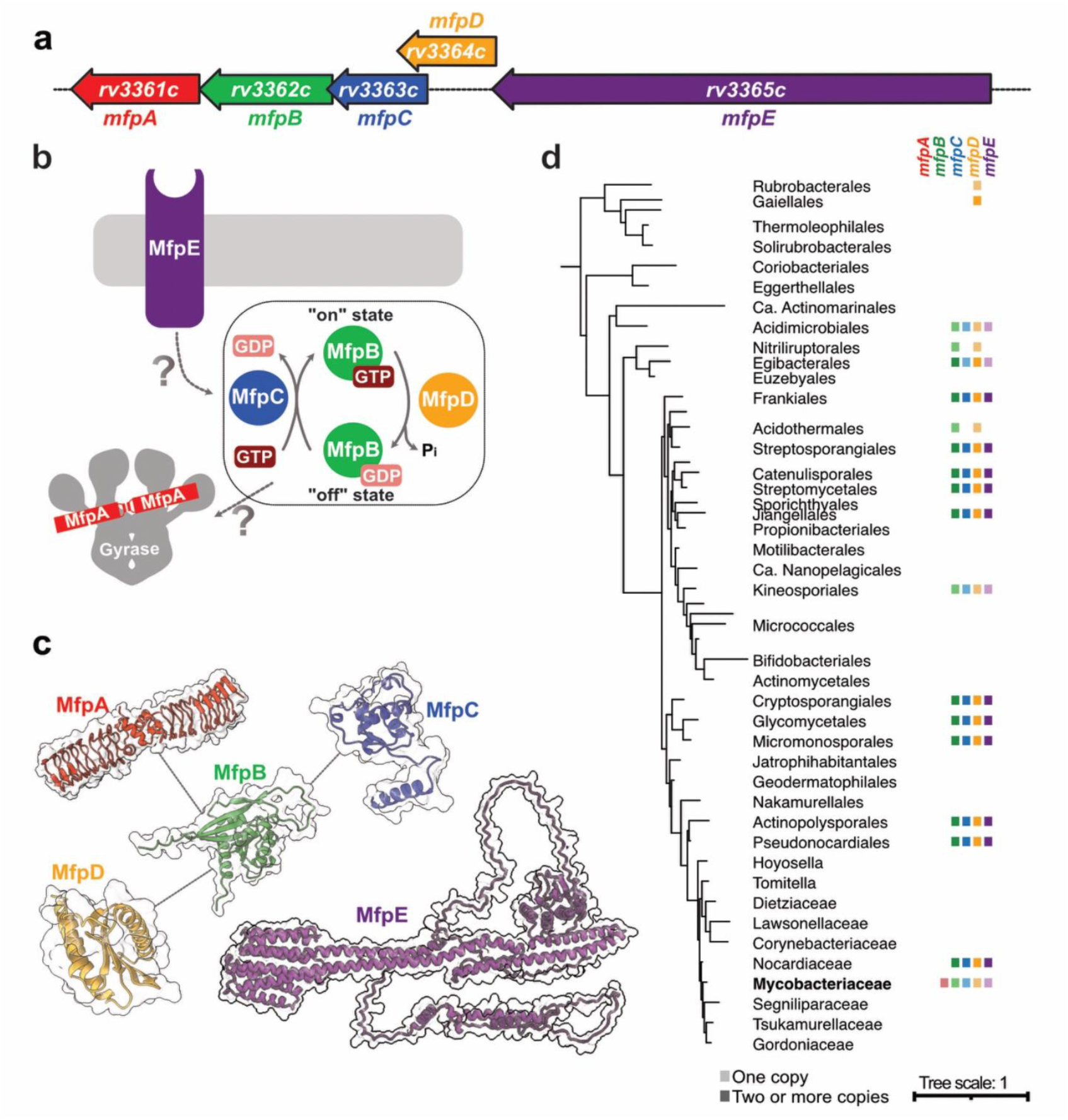
Organization, role and evolution of *Mycobacterium* FQ resistance proteins (Mfp). **a**, Operon organization from coding sequences rv3361c to rv3365c in *Mtb* genome. **b**, Proposed functional model of the five-membered Mfp team. MfpE is potentially a signal transducer that could alter the MfpB (GTPase)/MfpC (GAP)/MfpD (GEF) binary switch cycle, therefore impacting MfpA-mediated FQ resistance and providing DNA gyrase protection. **c**, Structural models of Mfp proteins. All structures have been generated in monomeric states with AF3 except for MfpA, which dimeric structures have already been reported (i.e. PDB 2BM6^18^). Validated protein-protein interactions between *Mtb* MfpA-MfpB^15^, *Msm* MfpB-MfpC^17^ and *Msm* MfpB-MfpD^16^ are represented in grey lines. **d**, Phyletic pattern of the presence of genes *mfpA*, *mfpB*, *mfpC*, *mpfD* and *mfpE* mapped onto a schematic reference phylogeny of the Actinobacteria. A colored square indicates the identification of the gene in at least half of the genomes analysed for the corresponding lineage. A darker shade of the color indicates the identification of more than one copy of the gene per genome. For the complete analysis, refer to Supplementary Figure 1.

Finally, previous studies show that MfpD also plays a distinctive role in pathogenicity during infection, making it of significant pharmacological interest^20,21^.

To date, from all the members of this five-component *mfp* system, only the *Mtb* and *Msm* orthologs of MfpA have been structurally characterized^19^. To gain further insight into this system, we chose to study the structural properties of *Mtb* MfpD using an integrative approach. Our initial findings from a phylogenetic analysis reveal that homologs of MfpB, MfpD and MfpE are widely present across most bacterial species. However, these proteins are found together with MfpC and MfpA as part of the five-membered *mfp* conservon exclusively within the Mycobacteriaceae family. Using X-ray crystallography, we demonstrated that MfpD shares structural similarities with other GAP proteins, particularly MglB of *Thermus thermophilus* (*Ttm*)^22^. By combining analytical size-exclusion chromatography and SAXS analysis, we confirmed that MfpD and MfpB can form a complex. Finally, by integrating these latter experimental results with previously published structural data on MglA/MglB homologs and AlphaFold3 models, we mapped the key interaction interfaces between MfpD and MfpB. This integrative analysis enabled us to propose a molecular mechanism underlying MfpB-mediated activation of the GTPase, and to demonstrate that MfpD forms a stable dimer in the presence of MfpB. Additionally, analysing the atomic structure of MfpD allowed us to identify the critical patch of residues involved in interactions with macrophage proteins. This confirms that these residues are ideally positioned and conserved exclusively within the Actinobacteria phylum.

## Results

### The *mfp* conservon has unique features in Mycobacteriacea

To determine the taxonomic distribution of the *mfp* conservon, we searched for homologs of its components in representative genomes from Actinobacteria. We found that *mfpB*, *mfpC*, *mfpD*, and *mfpE* are presents in almost half of the Actinobacteria genomes analyzed; however, *mfpA* was exclusively identified in the Mycobacteriaceae family of the order Mycobacteriales (*Figure 1d and Supplementary Figure 1*). In most cases, when these genes are present, they are organized in the genome as a conservon (*Supplementary Figure 2*). Interestingly, these conservons appear in multiple copies within each genome, except in Acidimicrobiales, Kineosporiales orders, some Nocardiaceae, and Mycobacteriaceae families, where only a single copy of each gene is found (*Supplementary Figure 2*).

Although widespread, the presence of these genes is scattered across the phylogeny of the phylum (*Figure 1d and Supplementary Figure 1*), suggesting the occurrence of duplications, losses, and/or horizontal gene transfer events during evolution. To better understand the evolution of these genes, we reconstructed a phylogeny of the *mfp* conservon by concatenating the protein sequences of each component (*Supplementary Figure 3*). The resulting phylogeny suggests that one copy of the conservon was likely inherited vertically during the evolution of Actinobacteria – which includes the single copy of the conservon in *Mtb* – and lost in some lineages, while the other copies appear to be the result of independent duplications and/or transfers (*Supplementary Figure 3*).

To further understand the origin of the *mfp* conservon, we searched for *mfp* genes in representative genomes from other bacterial phyla. We identified *mfpB*, *mfpD*, and *mfpE* genes in different bacterial phyla; however, *mfpC* is exclusively found in the Actinobacteria, and *mfpA* seems to have originated in the Mycobacteriaceae family. In the protein sequence alignment that included sequences from all bacteria, MfpB in Actinobacteria contains an insertion corresponding to positions 49 to 60 in *Mtb* (residues 70 to 80 on the alignment), which is absent in representative sequences from other bacterial phyla (*Supplementary Figure 4*). In the alignment of MfpD, we did not identify clear differences (*Supplementary Figure 5)*, while the alignment for protein MfpE showed a conserved N-terminal domain but a longer C-terminal domain in Actinobacteria, with no notable sequence conservation (*Supplementary Figure 6*).

### Biophysical properties and crystallization of MfpD

To get a structural insight into the organization of MfpD, we inserted the open reading frame *rv3364c* from *Mtb* H37Rv in pETM11 for expression with a histidine tag in N-terminal extremity. After overexpression in *E. coli*, a two-step purification procedure including a TEV protease-cleavage of the tag was conducted as detailed in the “Material and Methods” section. As shown with our SEC experiments using the purification buffer, MfpD eluted in three different peaks with peak (2) being the major one (*Figure 2a*). Protein samples from peak (2) were validated to be at the expected molecular weight for MfpD with SDS-PAGE. Protein quality was also assessed by dynamic light scattering (DLS), which revealed a major population peak with a hydration radius (RH) of 3.6 nm and a low polydispersity index (Figure 2b), and by nano-differential scanning fluorimetry (Nano-DSF), which revealed a melting temperature of 65°C *(Figure 2b, c*). These are favorable values for crystallization likelihood (*Figure 2c*)^23^ and reflect the compatibility of the buffer used to the best quality of the protein. To further investigate the hydrodynamic and oligomeric properties of MfpD, we performed SAXS and analytical ultracentrifugation (AUC) experiments (*Figure 2 d-g*). The SAXS scattering profile agrees with DLS results, showing a smooth decrease in intensity as a function of the scattering vector q, which is characteristic of monodisperse samples with no significant aggregation (*Figure 2d*). The dimensionless Kratky and P(r) plots also confirm that MfpD is a well folded, asymmetrical globular protein (*Figure 2e, f*). AUC measurements indicate that MfpD adopts a slightly elongated conformation, as reflected by the frictional ratio (*f/f0*) (*Figure 2g, h*). Both SAXS and AUC data consistently point to a dimeric state in solution, supported by experimental molecular weight estimations (*Figure 2h*). After HTS-crystallization trials, the hits were optimized and the best MfpD crystals were obtained in a solution consisting of 20 mM sodium acetate and 60% Methyl Pentanediol. MfpD crystallized in the P2(1) space group (*Table 1*) with two molecules in the asymmetric unit. The structure was solved at 1.8 Å resolution by molecular replacement using *Ttm* MglB (PDB ID: 3T1S) as a search model^22^, as identified through a DALI analysis.

**Figure 2.**
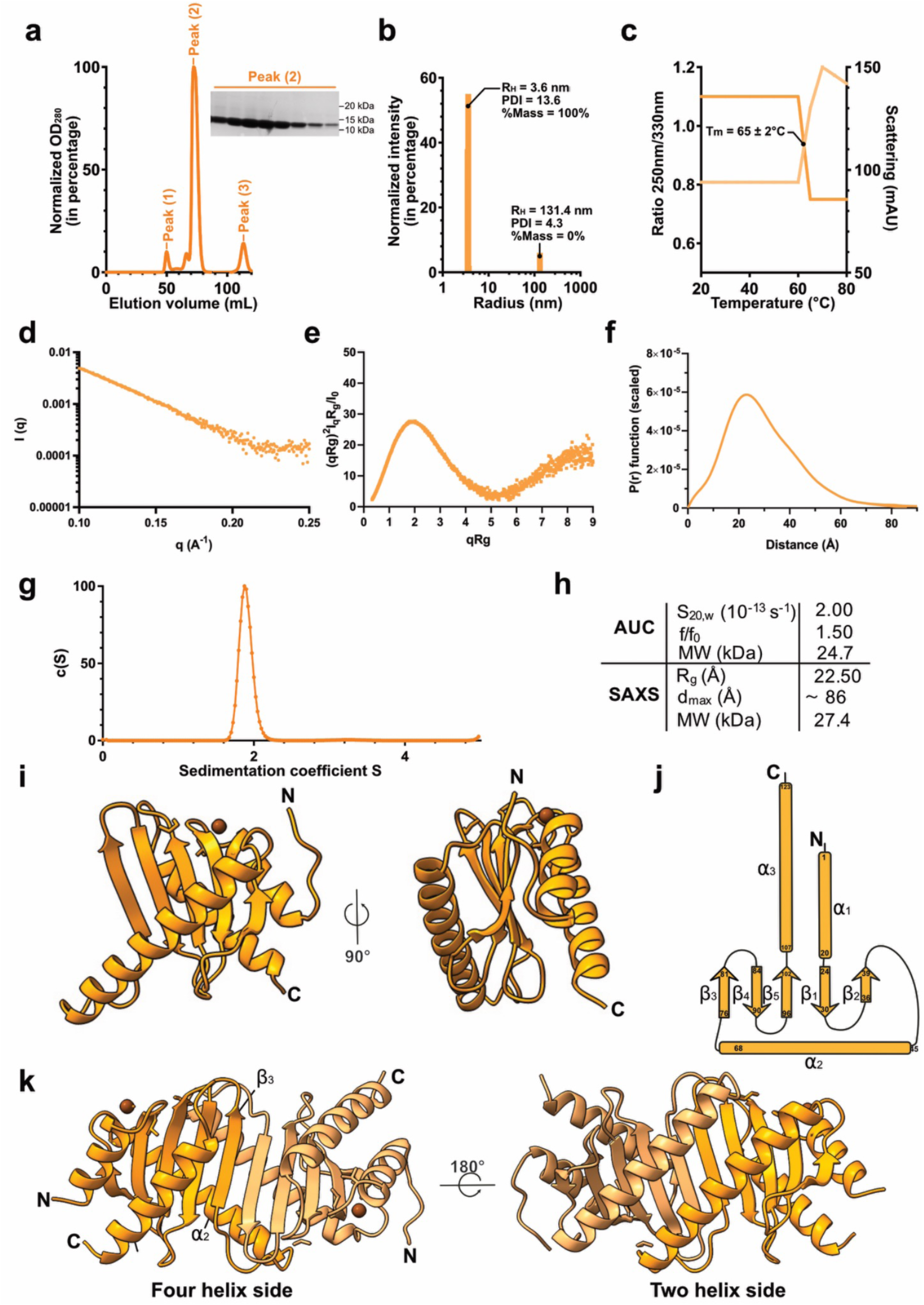
Biophysical and structural properties of *Mtb* MfpD. **a**, Size exclusion chromatography profile of purified *Mtb* MfpD on a Superdex 75 column and corresponding 8-12% SDS-PAGE for protein samples from peak (2). Molecular weight of wild-type *Mtb* MfpD without the histidine tag is equal to 13 kDa. **b-c**, Biophysical characterization of *Mtb* MfpD with (**b**) dynamic light scattering (DLS) and (**c**) nano-differential scanning fluorimetry (Nano-DSF) in optimized buffer A. **d-f**, SAXS data for MfpD shown as intensity curves (d), dimensionless Kratky plots (e) and P(r) plots (f). **g**, Analytical ultracentrifugation (AUC) continuous sedimentation coefficient c(S) distribution for 5 mg/mL of MfpD in the presence of 0.5 mM TCEP. Sedimentation coefficients are expressed in Svedberg units. **h**, Summary of hydrodynamic parameters for MfpD. **i**, Ribbon representation of *Mtb* MfpD monomer from crystal structure. **j**, Topological diagram of secondary structure of *Mtb* MfpD. **k**, Ribbon representation of *Mtb* MfpD dimer from crystal structure (f) with an emphasis on the dimerization interface.

**Table 1.** X-ray diffraction data collection and refinement statistics for Mtb MfpD (PDB 9HEF).

|  |  |
| --- | --- |
| **Synchrotron beamline** | SOLEIL PX2A |
| **Wavelength (Å)** | 0.98 |
| **Resolution range (Å) <sup>a</sup>** | 45.9 - 1.8 (1.9 - 1.8) |
| **Space group** | *P* 1 21 1 |
| **Unit cell** |  |
| ***a*, *b*, *c*** | 40.735, 73.65,<br>49.269 |
| **$\alpha$, $\beta$, $\gamma$** | 90, 111.38, 90 |
| **Total reflections** | 160519 |
| **Unique reflections** | 35875 (3487) |
| **Multiplicity** | 6.7 (2.0) |
| **Completeness (%)** | 99 (95.72) |
| **Mean *I*/sigma(*I*)** | 14.3 (1.3) |
| **Wilson B-factor** | 34 |
| **R-merge** | 0.028 (0.504) |
| **R-pim <sup>b</sup>** | 0.028 (0.504) |
| **CC1/2** | 99.9 (72.7) |
| **R-work (%) <sup>c</sup>** | 0.2324 (0.3740) |
| **R-free (%) <sup>c</sup>** | 0.2696 (0.3971) |
| **Number of non-hydrogen atoms** |  |
| **macromolecules <sup>d</sup>** | 1703 |
| **ligands** | 10 |
| **solvent** | 48 |
| **RMS deviations** |  |
| **Bond lengths (Å)** | 0.006 |
| **Bond angles (°)** | 0.76 |
| **Ramachandran favored (%)** | 99.58 |
| **Ramachandran allowed (%)** | 0.42 |
| **Ramachandran outliers (%)** | 0.00 |
| **Rotamer outlier (%)** | 0.60 |
| **Clashscore** | 1.16 |
| **Average B-factor** | 38.40 |
| **macromolecules** | 38.25 |
| **ligands** | 44.54 |
| **solvent** | 42.38 |
| **PDB ID** | 9HEF |

<sup>a</sup> Values in parentheses refer to the highest resolution shell.
<sup>b</sup> $R_{pim} = \sum_{hkl} [1/(N-1)]^{1/2} \sum_i |I_i(hkl) - \langle I \rangle(hkl)| / \sum_{hkl} \sum_i I_i(hkl)$ , where *N* is the multiplicity, *I<sub>i</sub>* is the intensity of reflection *i* and $\langle I \rangle(hkl)$ is the mean intensity of all symmetry-related reflections.
<sup>c</sup> $R_{work} = \sum ||F_o| - |F_c|| / \sum |F_o|$ , where *F<sub>o</sub>* and *F<sub>c</sub>* are the observed and calculated structure factor amplitudes. Five percent of the reflections were reserved for the calculation of *R<sub>free</sub>*.
<sup>d</sup> Values from MOLPROBITY<sup>37</sup>

### Overall crystal structure of MfpD

The MfpD monomer adopts an ɑ/β fold, an isomorphic structure that is present in other eukaryotic and prokaryotic proteins related to the Roadblock/LC7 family (Pfam PF03259) (*Supplementary Figure 7*). The (ɑββ)*2*βɑ architecture is organized as a central antiparallel 5-stranded β-sheets sandwiched between two ɑ-helices (*Figure 2i, j*). The protein adopts a globular shape with a calculated radius of gyration (R_g_) of 18.8 Å and a maximum particle dimension (d_max_) of 58.0 Å. MfpD monomers associate via the α2 and β3 regions, forming an extended antiparallel β-sheet that gives rise to a dimer characterized by two-helix and four-helix faces (*Figure 2k*). The dimerization interface between the two ɑ2 helices is predominantly stabilized by hydrophobic interactions involving residues Val54, Leu58, Leu61, Ala65, Leu68 and Phe69. In addition, hydrogen bonds reinforce the homodimer, mainly within the β-sheet region but also in the loops connecting α2 to β2 and β2 to β3 (*Supplementary Figure 8 and Supplementary Table 2*). The overall structure of each monomer is further stabilized via the presence of two Na*^+^* ions in the loop connecting ɑ1 and β1 secondary structures elements. Hydrogen bonds involving the main chains of Ala18, Val21, Val24 and one water molecule with the Na*^+^* ion of chain B, and Ala18, Val24, four water molecules, and the Na*^+^* ion of chain A, collectively contribute to the stabilization of the ɑ1 helix (*Supplementary Figure 9*). Additionally, no electron density corresponding to some regions of the N– and C-terminus regions (residues 1-8 and 125-130 in monomer A; and residues 1-2 and 126-130 in monomer B) has been observed in the map, suggesting their high flexibility.

### MfpD displays a GAP-like bacterial structure

For further comparative studies of *Mtb* MfpD, we considered the well-studied MglB from *Ttm*^22^. MglB, as the case for MfpD, contains a Roadblock/LC7 domain and has been shown to act as a GAP by activating the GTPase activity of MglA. MglB lacks the activating conserved residue (Arg, Asp or His) typically found in eukaryotic GAPs, and its MglA activation mechanism remains speculative. MglB also exists as a homodimer in crystal structures and in solution. First, a sequence comparison of the two proteins revealed 23% identity and 45% similarity (*Figure 3a*). However, superposition of the MglB and MfpD crystal structures shows r.m.s.d values of 1.0 Å for the monomer structures (83 atoms aligned, 3T1S) and 1.1 Å for the dimeric structures (77 atoms aligned, 3T1S). This indicates a similar overall fold, with slight differences in the relative positioning of the secondary structure elements. The main differences being found in the positions of the loops that connect the secondary structural elements in the N– and C-terminal extremities (*Figure 3b*). Additionally, according to a Consurf analysis, only a few primarily non-polar hydrophobic residues are evolutionarily conserved (*Supplementary Figure 8*). These residues are located at the contact interfaces on the surface of helix ɑ2 (*Figure 3c*). This helix is involved in the interaction with MglA in the case of MglB. Although these residues are not conserved the type of interactions that stabilizes the homodimer is conserved. Hydrogen bonds stabilize the β-strand sheet on one side of the homodimer, and hydrophobic interactions stabilize the interactions of the ɑ2 helix on the other side (*Supplementary Figure 8*). Additionally, we observed two sodium ions binding pockets in our MfpD crystal structure (*Supplementary Figure 9*).

**Figure 3.**
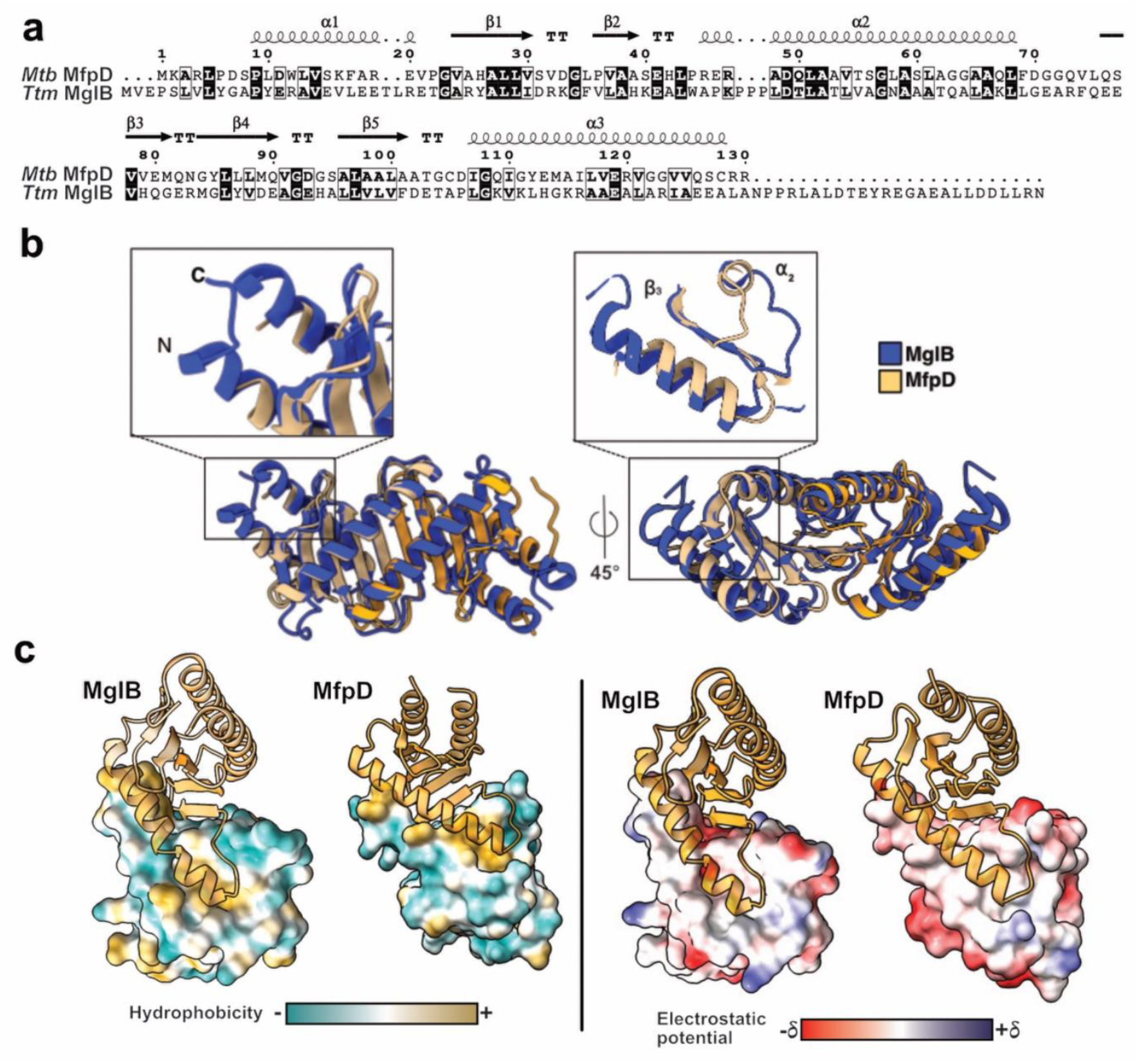
MfpD is homologous to MglB with distinctive evolutionary features. **a**, Alignment of *Mtb* MfpD and *Ttm* MglB. Conserved residues are highlighted in dark, similar residues are highlighted in boxes. The secondary elements of MfpD are shown above the alignment. **b**, Superposition of crystallographic structures of MfpD and MglB (present work and PDB:3T1S) dimers. **c**, Hydrophobicity and electrostatic potential comparison of the surface of *Mtb* MfpD and MglB.

### MfpB is homologous to MglA with a unique actinobacterial extended Switch I region

Small GTPases are diverse in sequence and catalytic mechanisms. Wuichet and Søgaard-Andersen^24^ previously proposed, after a detailed examination of the sequences of Ras superfamily of small GTPases in prokaryotes, a classification into two distinct families MglA and Rup with distinct catalytic mechanisms and suggested their involvement in numerous signaling pathways, including two-component systems. They also highlighted MfpB (termed MfpX) as being part of the MglA family. Our attempts to purify the full-length MfpB were unsuccessful. To address this issue, we generated AF3 models of MfpB in the absence or the presence of guanylate nucleotides. Firstly, we identified the five commonly conserved regions called the G domains (noted G1 to G5) and the essential regions named switch I, β2-screw and switch II on our structural MfpB model. Our analysis of the sequence and models of MfpB indicates that most of the conserved residues required for guanine nucleotide binding and GTP hydrolysis are conserved (*Figure 4a and Supplementary Figure 10*). The G1 region includes the P-loop, a signature structure in nucleotide-binding proteins that interacts with β-phosphate of GTP and magnesium ion, harboring the sequence *_21_*GXXXGKT*_27_* in MfpB. Moreover, we identified an extra-long switch I loop containing a unique Actinobacteria-specific insertion that connects β2 and β3 of MfpB (*Supplementary Figure 4*). This insertion in MfpB results in an extended loop, providing greater flexibility to this region. In the G2 region, also known as the β2-screw, one conserved threonine (Thr65), which interacts with ɣ-phosphate of GTP and magnesium ion, is present in MfpB. The consensus G3 motif, which typically follows the pattern DXXGQ/H/T and coordinates a water molecule acting as a nucleophile, is present as *_85_*GXXGQ*_89_* in MfpB. Therefore, the water molecule acting as a nucleophile is potentially coordinated in the active site by Gln89, similar to what is observed in MglA. The G4 region does not contain the [N/T]KxD motif – but instead features the *_137_*NEFD*_140_* motif, with Phe139 being implicated in ℼ-stacking with the guanine of the nucleotide. Finally, the G5 motif, which typically contains an essential serine that interacts specifically with guanine moiety of GTP, is not strictly conserved in MfpB, as it appears in the sequence *_166_*DARN*_169_*. In this case, the residues likely responsible for interacting with the guanine moiety are Arg168 (which may be involved in van der Waals interactions) and Asp166 (which could contribute to hydrogen bonding). The above analysis supports the classification of MfpB as a small GTPase related to the Ras superfamily. A comparison with its counterpart in Ttm, MglA, shows that the two enzymes are indeed highly similar (*Figure 4a-e*).

**Figure 4.**
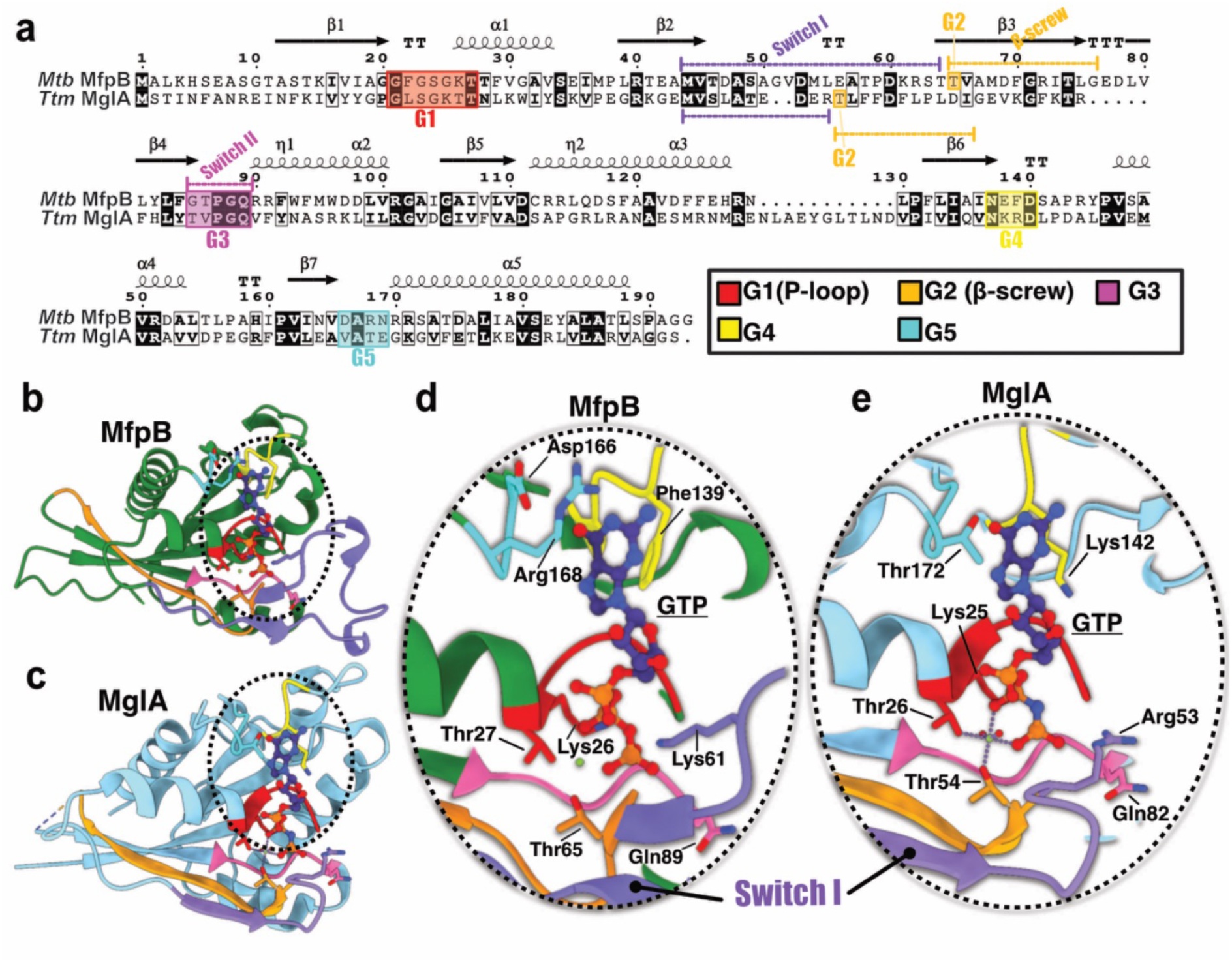
Structural comparison of MfpB versus MglA. **a**, Alignment of *Mtb* MfpB and *Ttm* MglA. The G1-G5 motifs and switch regions characteristic for the G domain and the secondary elements of MfpD are shown below and above the alignment, respectively. **b-c**, AF3 model of MfpB (b) and crystallographic structure of MglA (PDB:3T12) (c). G1-G5 motifs and switch regions are also highlighted as in **a**. **d-e**, Zoom on the GTP binding sites.

### Molecular and structural insight into the MfpB-MfpD interaction

MfpB and MfpD activities have been previously validated to be functional *in vitro*, with MfpD being a GAP for MfpB^17^, but little is known about their structural organization. We thus co-expressed the two *Mtb* proteins from a pET-DUET system in *E. coli*, and co-eluted both proteins thanks to the presence of a histidine tag only on MfpD, with the complex remaining formed after SEC. As shown in *Figure 5a*, SEC indicates that the MfpB/MfpD proteins form a stable and non-transient complex. Furthermore, the interaction with MfpB stabilizes the MfpD dimer, making it sufficiently stable to resist denaturation under SDS-PAGE conditions (*Figure 5a)*. This supports the idea that co-expression of MfpD and MfpB promotes the solubility of MfpB under our experimental conditions. Activity assays on the complex MfpB/MfpD enabled us to detect a GTPase activity, meaning that MfpB within the complex is functionally active, with a basic optimal pH at 9-10 (*Figure 5b*).Then, we analyzed this complex with SAXS experiments in the absence or the presence of 5 mM of GDP or non-hydrolysable GTP analogue, MgGTP*γ*S *(Figure 5 c-e)*. Intensity scattering curves firstly show similar profiles in all three conditions. A slight peak shift in the presence of MgGTP*γ*S is observed, meaning that slight conformational changes occur after binding of GTP. Nonetheless, no differences are noted in apo form or in the presence of GDP (*Figure 5c*). Further inspection of the dimensionless Kratky plots shows Gaussian curves in all tested conditions, meaning that MfpB-MfpD form a well-folded globular complex *(Figure 5d)*. P(r) functions lead us to identify an induction of a slight rightward shift only in the presence of MgGTP*γ*S, meaning that low protein dimension expansion occurs after binding of these substrates *(Figure 5e).* Mass photometry experiments for MfpB-MfpD concordantly showed the existence of a singular population corresponding to a molecular weight of 53 kDa, meaning that the complex is formed by two monomers of MfpD and one monomer of MfpB (*Figure 5f*).

**Figure 5.**
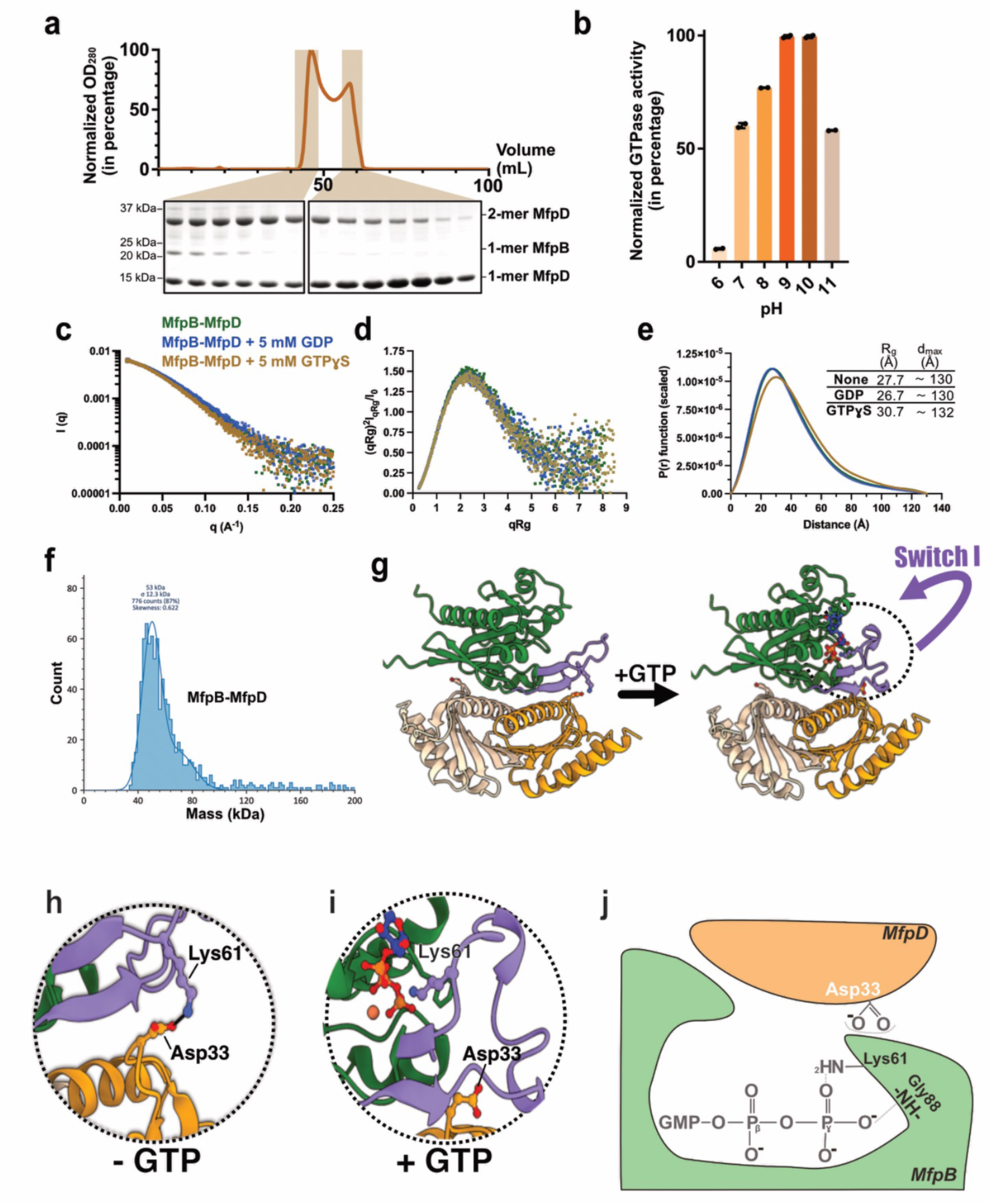
*Mtb* MfpB-MfpD interaction. **a**, SEC of MfpB-MfpD complexes (top) and 4-12% SDS-PAGE analysis of proteins from the two major peaks (bottom). **b**, GTPase activity of MfpB from the MfpB-MfpD mixture in different pH conditions. **c-e**, SAXS data for MfpB-MfpD in the absence (green) or the presence of 5 mM GDP (blue) or GTP (brown) showing the comparisons of intensity curves (c), dimensionless Kratky plots (d) and P(r) plots. **f**, Mass photometry profiles for 20 nM of co-eluted MfpB-MfpD. The peak corresponds to a molecular weight of 53 kDa. **g-i**, AF3 models of 1MfpB-2MfpD complexes in the absence or presence of GTP (f) and zoom on corresponding switch I regions (**h** and **i**). **j**, Proposed mechanism for GTP binding in the case of MfpB and MfpD.

### MfpD may function as a GAP for MfpB

Similarly to MglB proteins, MfpD lacks the conserved arginine, asparagine or histidine residue that is involved in stimulating GTP hydrolysis in Ras proteins, making it unable to catalyze GTP hydrolysis without the intervention of residues from MglB. Indeed, in the case of MglA, it has been previously shown that conformational changes upon interaction with MglB induce repositioning of important residues in the active site for GTP hydrolysis^22^ (*Figure 4*). This includes a highly conserved arginine (Arg53) that moves into a position where it contacts the ɣ-phosphate of GTP, as well as a conserved glutamine (Gln82) that positions itself towards the ɣ-phosphate and the nucleophilic water molecule (*Figure 4e*).

Therefore, we compared the sequences and the AF3 model for a MfpB:MfpD (1:2 stoichiometry) complex with the equivalent *Ttm* MglA:MglB heterotrimer. This analysis revealed that the arginine 53 is not conserved in actinobacteria GTPases (*Supplementary Figure 4*). Inspection of the model and the sequence alignment shows that a highly conserved lysine (Lys61) could potentially play the role of the arginine, although the efficacy of hydrolysis might be reduced (*Figure 5 g-j, Supplementary Figure 11*).

Moreover, a previous study also highlighted the potential involvement of a conserved negatively charged residue from the GAP protein located in its loop between strands β1 and β2 that is positioned directly in the active site after GTPase-GAP interaction^18^. This residue is a lysine in MglB (Lys38) and an aspartate in MfpD (Asp33). Upon examining the position of this Asp33 in the crystallographic structure of MfpD, we observed that it is located on the surface of the protein within the loop connecting strands β1 and β2 (*Figure 5g and Supplementary Figure 12*). Analysis of the AF3 model of the 1MfpB-2MfpD heterotrimer indicates that this residue is positioned at the contact interface between MfpD and MfpB. This positioning is consistent with an important role in activating GTPase hydrolysis by inducing the essential conformational changes needed to position Lys61 correctly in the active site (*Figure 5g-j*). A comparison of two AF3 models of the heterotrimer in apo form or in the presence of GDP or GTP reveals a major conformational change of the switch I loop. When GTP is bound, the switch I loop bends inward toward GTP-binding site, positioning the Lys61 side chain toward the ɣ-phosphate of GTP (*Figure 5i*). In contrast, in the absence of GTP, the switch I loop adopts an extended conformation, with Lys61 being oriented toward the MfpD interface, where it forms a salt bridge with Asp33 (*Figure 5h*). This interaction may contribute to the stabilization of this heterotrimer in the absence of GTP. This may also explain the increased efficiency of GTP hydrolysis at higher buffer pH. As the pH rises, lysine becomes progressively deprotonated, weakening the salt bridge and facilitating the conformational change of the switch I loop.

Finally, this heterotrimer is further stabilized via several hydrogen bonds linked to the presence of the Asp33 from the other MfpD chain of the MfpD dimer, inside a very electropositive pocket inside MfpB and hydrophobic interactions between the two ɑ2 helices of the MfpD dimer and MfpB (*Supplementary Figure 13)*.

## Discussion

Many bacterial species have evolved sophisticated regulatory systems involving GTPases to modulate diverse cellular processes, such as metabolism, cell motility and polarity, and pathogenicity^25,26^. In *Mtb*, the *mfp* conservon, linked to FQ resistance in *Mtb*, comprises a conserved triplet MfpB (a GTPase), MfpC (a putative GEF), and MfpD (a GAP protein). This conserved signaling unit appears broadly distributed across bacteria. At the functional level, MfpD was previously identified as an activator of MfpB’s catalytic activity, triggering its transition from an active “on” to an inactive “off” state. Moreover, MfpD is of supplemental interest because it has been shown, in the context of an infection in the presence of macrophages, to be secreted from mycobacteria and to play an essential role in evading host defense mechanisms by binding to human membrane-associated serine protease cathepsin G protein^27^.

In this study, we focused on the structural and biophysical characterization of *Mtb* MfpD and its interaction with its cognate GTPase MfpB. Our approach, combining AUC, SAXS, and X-ray crystallography, revealed that MfpD adopts a nearly globular shape and a dimeric organisation in solution. The crystal structure shows that MfpD adopts a α/β fold, characteristic of the Roadblock/LC7 protein family^28^, with notable similarity to *Ttm* MglB, despite limited sequence identity. This family of proteins are linked to proteins involved in motility in eukaryotes, such as the flagellar outer arm or cytoplasmic dynein^29^, as well as polarity regulation and gliding motility in bacteria (e.g. *Myxococcus xanthus*^26,30–32)^. The dimeric form is stabilized by hydrophobic interactions within the α2 helices. The oligomeric state of MfpD appears to be a conserved architecture as the case of MglB in MglA-MglB, despite the variability in residues implicated in dimerization. Dimerization may represent an intricate strategy for spatial control of GTPase signaling in bacteria.

We modeled the structure of MfpB, previously characterized as a Ras-like GTPase, and compared it to MglA from *Ttm*. This analysis enabled the identification of conserved G-domain motifs critical for GTP binding and hydrolysis. A particularly striking feature of MfpB is the extended switch I region, unique to Actinobacteria, which may confer additional conformational flexibility and functional specificity. From a mechanistic standpoint, our MfpB-MfpD model proposes that MfpD enhances GTP hydrolysis activity of MfpB through conformational stabilization rather than a direct catalytic residue insertion, similarly to the atypical GAP mechanism observed for MglB-MglA. The conservation of the Asp33-Gly34 motif across Actinobacteria also suggests that this local spot may have co-evolved to substitute the canonical arginine found in eukaryotic Ras-GAP proteins, and thereby fine-tuning the catalytic efficiency of MfpB within the unique metabolic and structural constraints in Mycobacteria.

Beyond its intracellular role in bacteria, MfpD has already been the basis of bioinspiration by Lee and colleagues for the conception of small molecules acting as antibacterial and anti-inflammatory agents. Indeed, after their description of MfpD as being implicated in interactions with host proteins SNX9 and YEATS4, the authors reported peptide-mimetic molecules based on the pharmacophoric features at the helical N-terminal sequence (*_12_*WLVSKF*_17_*) of MfpD, identified as being at the interaction interface with SNX9 and YEATS4^33^. Linked to these observations, and by inspecting MfpD sequence conservation, we observed a high conservation of the hydrophobic nonpolar amino acids Trp12 and Phe17 across species (*Supplementary Figure 5*). Structurally, this motif is positioned within the ɑ1 helix of MfpD, with the side chain of Trp12 extended to the solvent (*Supplementary Figure 14*). The conserved motif in MfpD is located on the opposite side to the MfpB-MfpD interaction interface, suggesting that MfpD could plausibly perform both functions independently. The consistent localization of Trp12 at potential interaction interfaces underscores the need for further structural studies.

More generally, the physiopathological relevance of the mfp conservon in *Mtb* remains incompletely understood and future work on the elucidation of the dynamic cycle of Mfp proteins is of interest. In *in vitro* conditions, all Mfp proteins are non-essential for growth as shown by phenotypic profiling and high density mutagenesis analysis in *Mtb*^20,21,34^. Moreover, high-scale systematic bacterial two-hybrid experiments have revealed that MfpB may also interact with DNA binding proteins or metabolic enzymes in *Mtb*^34^. RNA-sequencing experiments on *Mtb* treated with kinases inhibitors also showed a significant decrease in mfpD in *Mtb*^35^. More intriguingly, through a genome-wide regulator-DNA interaction one-hybrid approach, all five Mfp proteins have been proposed to be part of the transcriptional regulatory network in *Mtb*^36^.

In summary, our study bridges the previous functional knowledge on MfpD and MfpB with structural and biophysical observations. The insights presented here may serve as a framework for future studies exploring both the fundamental biology and therapeutic targeting of this signaling system.

## Methods

### Sequence analyses

To study the taxonomic distribution of MfpA, MfpB, MfpC, MfpD and MfpE in Actinobacteria phylum, we performed sensitive HMM similarity searches against a database containing 244 taxa representing all Actinobacteria diversity^38^. We used the HMMER package (v.3.3.2)^39^ tool ‘jackhmmer’ to look for homologues of the proteins coded by the *mfp* genes in all the proteomes, using the GenBank^40^ sequences Rv3361c, Rv3362c, Rv3363c, Rv3364c and Rv3365c as queries. The hits were aligned with mafft (v.7.475) using default parameters. Alignments were manually curated, removing sequences that did not align. The hits obtained by jackhmmer might not include sequences that are very divergent from the single sequence query. For this, the curated alignments were used to create HMM profiles using the HMMER package tool ‘hmmbuild’. Curated HMM profiles for each protein were used for a final round of searches against the Actinobacteria database, using the HMMER tool ‘hmmsearch’. All hits were curated to remove false positives by checking alignments obtained using linsi, the accurate option of mafft (v.7.475)^41^. Finally, we analysed the taxonomic distribution of the identified proteins. The phyletic pattern and the genomic context information were mapped on an Actinobacteria reference phylogeny using iTOL^40^. We used the same HMM profiles to look for the Mfp proteins in a database containing 81 representative genomes of all Bacteria phyla (five taxa per phylum with cultured representatives)^42^. To reconstruct the *mfp* conservon phylogeny, we concatenated the protein sequences of the proteins coded by each conservon. Homologues of these proteins were aligned as explained before, and trimmed using trimAl^43^, keeping the columns that contain less than 20% of gaps. We used this alignment to reconstruct a maximum-likelihood phylogeny with IQ-TREE^42^ using the Model Finder Plus (MFP) option, with Ultrafast Bootstrap supports calculated from 10.000 replicates, with a minimum correlation coefficient of 0.999.

### Cloning

pETM11-MfpD plasmid constructed with isothermal Gibson assembly^44^ harbors the open reading frame (ORF) *rv3364c* of MfpD of *Mtb* and enables the recombinant expression of MfpD with a histidine tag in the N-terminus. In brief, *rv3364c* was amplified from M. tuberculosis H37Rv genome with forward (5’-AAGAAGGAGATATACATATGGGTAAGGCGCGCGTCTGCCG-3’) and reverse (5’-TTACCAGACTCGAGGGTACCTTAACGACGGCAGCTTTGAACC-3’) primers (Eurofins MWG Operon) with Phusion® High-Fidelity DNA polymerase (New England Biolabs). PCR products were extracted by gel electrophoresis (0.8% agarose in Tris-acetate-EDTA buffer) and purified by NucleoSpin Gel and PCR Clean-up kit (Macherey-Nagel). Gibson assembly was conducted with Gibson Assembly® Master Mix (New England Biolabs) as recommended by the manufacturer. After transformation in E. coli DH5ɑ strain, clones were selected and plasmid isolation was done using QIAprep miniprep kit (Qiagen). The sequence was verified by Sanger sequencing (Eurofins Genomics).

pDUET-MfpB plasmid purchased from GenScript Biotech harbors the ORF *rv3362c* of MfpB of M. tuberculosis H37Rv and enables the recombinant expression of a tag-free MfpB.

### Overexpression and purification of Mtb MfpD

*E. coli* Bli5 (DE3) strain was transformed with a pETM11-MfpD construct. The protein was produced by adding 0.5 mM of isopropyl-β-D-thiogalactopyranoside (IPTG) during the exponential phase of growth. After 16h at 30°C, cells were harvested by centrifugation and resuspended in 20 mM Tris-HCl, pH 8, 500 mM NaCl, 10 mM imidazole, 5 mM MgCl*_2_*, 1 mM DTT, and 10% (v/v) glycerol. After sonication with a Bioblock Scientific Vibra Cell, the mixture was supplemented with a cOmplete*^TM^*Mini EDTA-free tablet (EASYpack, Roche Diagnostics) and centrifuged. The soluble extract was loaded on a Ni*^2+^*-affinity column (HisTrap HP column, Cytiva) and the His-tagged protein was purified by applying a linear imidazole gradient using a buffer containing 20 mM Tris-HCl, pH8, 500 mM NaCl, 250 mM imidazole, 5 mM MgCl*_2_*, 1 mM DTT, and 10% (v/v) glycerol. Fractions of interest, identified by SDS-PAGE^45^, were pooled and subjected to TEV digestion for 18 hours at 4°C in buffer A (20 mM Tris-HCl, pH 8.5, 300 mM NaCl, 1 mM DTT, 1 mM MgCl*_2_* and 5% glycerol). After a separation on a Ni*^2+^*-affinity column, the flow-through was collected, concentrated and further purified by size-exclusion chromatography (Superdex 75 column, Cytiva) equilibrated in buffer A. Proteins were flash-frozen in liquid nitrogen and stored at – 80°C. Protein concentration was determined by using the molar absorption coefficient predicted from the amino acid sequence by the ProtParam tool (http://web.expasy.org/protparam/)^46^.

### Co-expression and co-elution of Mtb MfpB-MfpD

*E. coli* Bli5 (DE3) strain was co-transformed with pETM11-MfpD and pDUET-MfpB. Proteins were produced through lactose-driven autoinduction. After 16h at 30°C, cells were harvested by centrifugation and resuspended in 20 mM Tris-HCl, pH 8, 200 mM NaCl, 0.1% (v/v) Triton and 10 mM imidazole supplemented with a cOmplete™ ULTRA EDTA-free protease inhibitor tablet (Roche Diagnostics). After bacterial lysis with a Cell Disruption Lysis System, the soluble extract was added to a His-Tag cOmplete*^TM^* metal affinity resin (Roche) and proteins were purified as recommended by manufacturer by using a buffer containing 20 mM Tris-HCl, pH8, 200 mM NaCl, 0.1% (v/v) Triton and 500 mM imidazole. Fractions of interest, identified by SDS-PAGE, were pooled and concentrated. Further purification by size-exclusion chromatography (Superdex 75 column, Cytiva) equilibrated in buffer A (50 mM Tris-HCl pH8, 50 mM NaCl, 1 mM MgCl*_2_*, 1 mM DTT, 5% (v/v) glycerol) was conducted. Further final steps were conducted as the case of MfpD purification.

### Dynamic light scattering (DLS) measurement

The homogeneity of the sample was verified at 20°C by DLS experiments on a DynaPro Plate Reader III (Wyatt technology) with a wavelength of 658 nm. Experiments were conducted on protein samples at 1 mg/mL loaded into an adapted 384-well microplate. Data were collected at a speed of ten acquisitions of 5 seconds each on the DYNAMICS version V.7.10.0.21 software. The selection of the buffer was conducted on protein samples with several buffer conditions as previously described^47^.

### Nano-differential scanning fluorimetry (Nano-DSF) experiments

Nano-DSF experiments were conducted with 10 μL protein samples at 0.1 mg/mL charged into a Prometheus NT.48 Series Nano-DSF Grade High Sensitivity capillary tube and onto a Prometheus NT.48 (Nanotemper) instrument. Tryptophan fluorescence emission was monitored at 330 nm and 350 nm. Measurement of the fluorescence emission was conducted in buffer C on a ramp of temperatures ranging from 20°C to 95°C. Data was processed with the Nanotemper Software.

### Analytical ultracentrifugation (AUC)

AUC experiments were carried out using a Beckman-Coulter ProteomeLab XL-I. MfpD was diluted in its corresponding buffer A at 5 mg/mL supplemented with 0.5 mM tris(2-carboxyethyl)phosphine (TCEP). Samples were centrifuged at 42.000 rpm in an 8-hole An-60 Ti rotor equipped with 1.2-mm thick two-channels epon-aluminium centerpiece with sapphire windows. Measurements of protein concentration as a function of the radial position and time were performed at 20°C by absorbance detection. Sedimentation velocity data analysis was then performed by applying a continuous size distribution analysis c(S) using Sedfit 18.1 software^48^. Frictional ratios (f/f*0*) were calculated from the sedimentation coefficient and the molecular mass according to the Svedberg equation*^49^*c. Protein partial specific volume from the amino acids sequence of MfpD (<u>ρ</u>*_i_* = 0.729 mL/g), and buffer A density (*ρ =* 0.0110 g/mL) and viscosity (*η =* 0.0103 poise) were estimated with SEDNTERP software^50^ (available from the Boston Biomedical Research Institute).

### Mass photometry (MP)

MP experiments were performed on a Refeyn TwoMP instrument precalibrated with bovine serum albumin and urease according to the manufacturer’s protocol. Protein samples were diluted to a final concentration of 20 nM in 50 mM Tris-HCl pH8. Data were collected for one minute with the software AcquireMP (Refeyn) and analyzed with the software DiscoverMP (Refeyn).

### Size exclusion chromatography small-angle X-ray scattering (SAXS)

X– ray scattering was recorded on MfpD and MfpB-MfpD after elution of 5 mg/mL of protein in the presence or absence or GDP or GTP on Superose™ 6 Increase 5/150 GL size exclusion chromatography column (Cytiva) on the SWING beamline at Synchrotron SOLEIL (Saint-Aubin, France) with a CCD-base AVIEX detector. The X-ray wavelength was 1Å and the beam geometry was 0.8 mm x 0.15 mm. X-ray exposure time was set at 1 second. Corresponding scattering vectors were processed as previously described^51^. Subtractions for intensity curves were conducted with FOXTROT. Data were processed and analyzed using the program PRIMUS^52^ from ATSAS 3.0.5 software suite^53^.

### Crystallization and data collection

Crystallization screening trials were carried out by the vapor diffusion method using a Mosquito TM nanodispensing system (STPLabtech, Melbourn, UK) following established protocols^54^ with purified MfpD (14mg/ml). Initial hits were reproduced by hanging-drop vapor diffusion at 18 °C.

The best MfpD crystals were grown from a solution consisting of 20 mM Na Acetate and 60 % MPD. Before cryocooling, single crystals were soaked in reservoir solution containing up to 30 %(w/v) ethylene glycol. The higher resolution X-ray diffraction data were collected on PROXIMA 2 beamline at SOLEIL Facility, Saclay, France using a wavelength of 0.98 Å (Table 1).

### Structure determination and refinement

Data processing was carried out using XDS^55^ and AIMLESS (which is part of the CCP4 software package^56^. Structure determination was performed by molecular replacement using 3T1S as the search model^22^. Two molecules in the asymmetric unit were observed at 1.8 *Å* resolution. The obtained solution underwent rigid-body refinement using PHENIX^57^. The density map was examined in Coot^58^ and the model was built using Coot and iterative rounds of refinements in PHENIX^57^ at the end, including the use of TLS parameters^59^. For PHENIX, before each refinement, the model was prepared using READYSET^57^. The final refinement statistics are reported in Table 1. All molecular images were generated using ChimeraX.

### GTPase activity assay

Hydrolysis of GTP was monitored according to manufacturer’s recommendations on a Sunrise plate reader (TECAN) with the ATPase/GTPase activity assay kit (Sigma, MAK113), based on the measurement of the absorbance of complexed malachite green with free phosphate at 620 nm.

### In silico model building with AlphaFold

Structural predictions were performed using the AlphaFold Server BETA powered by AlphaFold3^60^. To ensure unbiased predictions for the different protein forms, structural templates were disabled in these cases. For the prediction of the Mfp complexes, all models converge to similar conformations and only the best one for each run is shown. The predicted template modeling (pTM) score and the interface predicted template modeling (ipTM) score for each structure^60–62^ are reported in *Supplementary Table 4*.

### Software

Computational analyses were performed using the SBGrid software environment, which provides curated, version-controlled structural biology applications and reproducible execution environments across platforms^63^.

### Data availability

Atomic coordinates and structure factors for MfpD have been deposited in the Protein Data Bank with accession number 9HEF (*Table 1*). Source Data for sequence analysis will be available in a Mendeley repository under accession code 10.17632/pd2yg9z226.1 upon publication, but is temporarily accessible from: https://data.mendeley.com/preview/d6hmm7bjnt?a=9155e04d-88b1-4719-8ec6-0a300aa479f6. Other inquiries are available from the corresponding author upon request.

## Acknowledgments

This work was supported by the IdEx Université Paris Cité, ANR-18-IDEX-0001 (to S.P.), the Centre National de la Recherche Scientifique (CNRS), Université Paris Cité and Institut Pasteur. Molecular graphics were done with ChimeraX, developed at UCSF with support from NIH (R01-GM129325) and NIAID. The authors would like to thank the staff of the Institut Pasteur “Plate-Forme de Cristallographie” for robot-driven crystallization screening. We thank the beamline scientists from the Synchrotron SOLEIL (St. Aubin, France) for advice in X-ray diffraction data collection on PROXIMA-2 beamline, and SAXS data collection on SWING beamline.

## Author contributions

A.G. and S.P. conceptualized and designed the project; S.P. supervised the work; A.G., M.M., D.M., E.L., E.B., B.R., A.H. and S.P. performed the experiments; A.G, M.M., D.M., B.R. and S.P. analyzed the data; P.M.A. and C.M. provided scientific guidance; A.G., D.M. and S.P. wrote the article with input from all authors.

## Competing interest

The authors declare no competing interests.

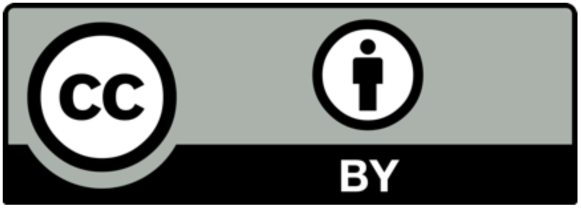

## Supplementary Information

**Supplementary Table 1.**
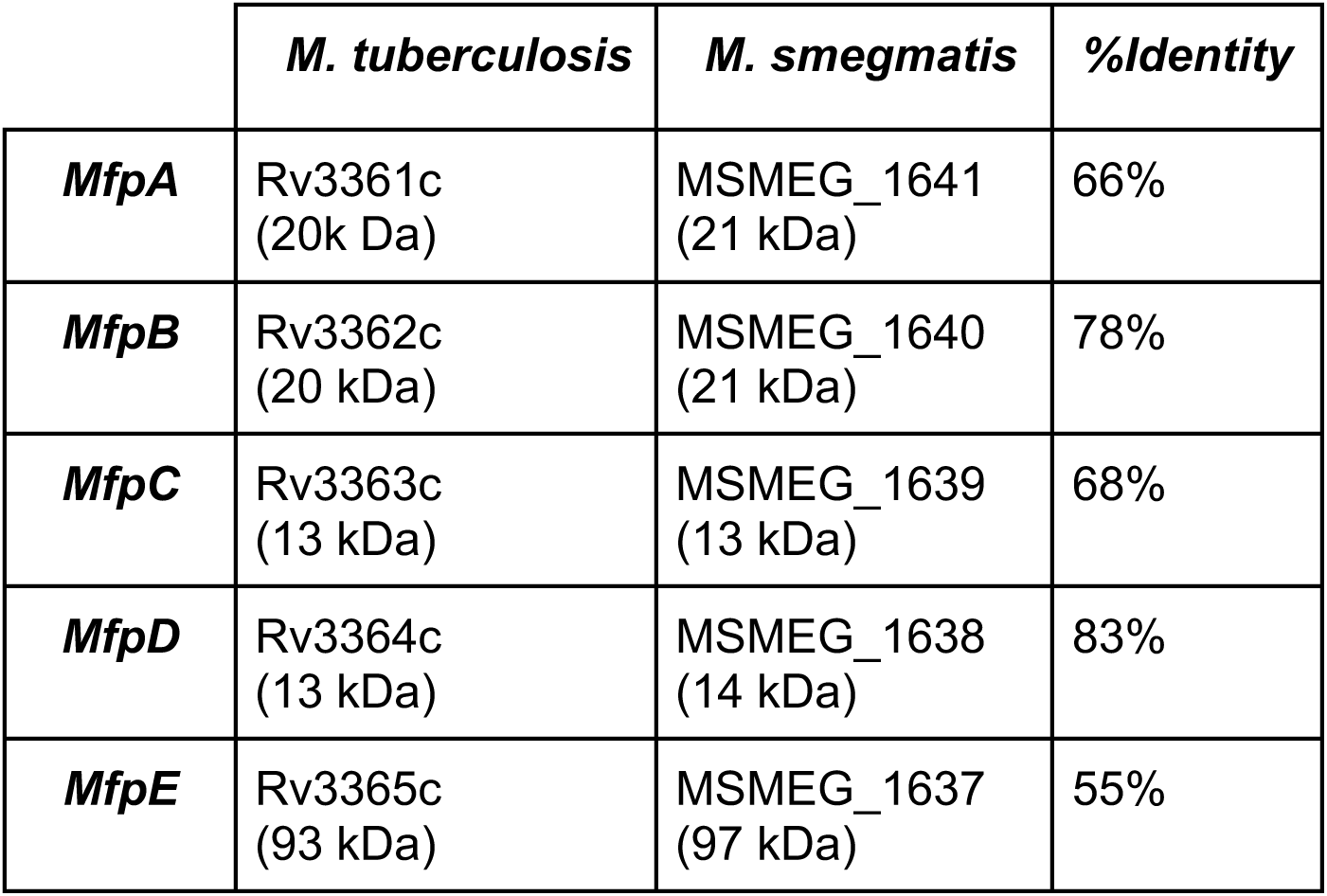
Identity percentages for Mfp proteins between M. tuberculosis H37Rv and M. smegmatis ATCC 700084. Molecular weights are listed for reference.

**Supplementary Table 2.** Hydrogen bonds implicated in MfpD dimerisation. Interactions are detected according to a PISA analysis. Note that no disulfide bonds, covalent bonds, or salt bridges exist.

|  | **Structure 1** |  | **Distance (Å)** | **Structure 2** |  |
| --- | --- | --- | --- | --- | --- |
| **1** | B:Arg | 47 (NH1) | 3.01 | A:Leu | 68 (O) |
| **2** | B:Gln | 73 (N) | 2.92 | A:Gln | 82 (OE1) |
| **3** | B:Leu | 75 (N) | 2.81 | A:Glu | 80 (O) |
| **4** | B:Gln | 76 (N) | 3.35 | A:Glu | 80 (O) |
| **5** | B:Gln | 76 (NE2) | 2.72 | A:Glu | 80 (OE1) |
| **6** | B:Val | 78 (N) | 2.99 | A:Val | 78 (O) |
| **7** | B:Glu | 80 (N) | 2.81 | A:Gln | 76 (O) |
| **8** | B:Gln | 82 (N) | 3.22 | A:Gly | 71 (O) |
| **9** | B:Gln | 82 (N) | 2.90 | A:Gln | 73 (O) |
| **10** | B:Asn | 83 (N) | 2.93 | A:Gly | 71 (O) |
| **11** | B:Asn | 83 (ND2) | 3.04 | A:Asp | 70 (O) |
| **12** | B:Leu | 68 (O) | 2.91 | A:Arg | 47 (NH1) |
| **13** | B:Gln | 82 (OE1) | 3.00 | A:Gln | 73 (N) |
| **14** | B:Glu | 80 (O) | 2.77 | A:Leu | 75 (N) |
| **15** | B:Glu | 80 (O) | 3.27 | A:Gln | 76 (N) |
| **16** | B:Val | 78 (O) | 2.95 | A:Val | 78 (N) |
| **17** | B:Gln | 76 (O) | 2.82 | A:Glu | 80 (N) |
| **18** | B:Gly | 71 (O) | 3.32 | A:Gln | 82 (N) |
| **19** | B:Gln | 73 (O) | 2.91 | A:Gln | 82 (N) |
| **20** | B:Gly | 71 (O) | 2.80 | A:Asn | 83 (N) |
| **21** | B:Asp | 70 (O) | 3.03 | A:Asn | 83 (ND2) |

**Supplementary Table 3.** Hydrogen bonds (entries 1 to 16) and salt bridges (entries 17-18) implicated in Ttm MglB dimerisation. Interactions are detected according to a PISA analysis. Note that no disulfide bonds or covalent bonds exist.

|  | **Structure 1** |  | **Distance (Å)** | **Structure 2** |  |
| --- | --- | --- | --- | --- | --- |
| **1** | A:Gln | 83 (N) | 2.71 | A:Gln | 88 (O) |
| **2** | A:Glu | 84 (N) | 3.18 | A:Gln | 88 (O) |
| **3** | A:Val | 86 (N) | 2.89 | A:Val | 86 (O) |
| **4** | A:Gln | 88 (N) | 2.84 | A:Glu | 84 (O) |
| **5** | A:Glu | 90 (N) | 3.71 | A:Glu | 79 (OE1) |
| **6** | A:Arg | 91 (NH1) | 3.10 | A:Leu | 77 (O) |
| **7** | A:Arg | 91 (NH2) | 2.31 | A:Gly | 78 (O) |
| **8** | A:Met | 92 (N) | 3.42 | A:Glu | 79 (OE1) |
| **9** | A:Leu | 77(O) | 3.70 | A:Arg | 91 (NH1) |
| **10** | A:Gly | 78 (O) | 2.31 | A:Arg | 91 (NH2) |
| **11** | A:Glu | 79 (OE1) | 3.42 | A:Met | 92 (N) |
| **12** | A:Glu | 79 (OE1) | 3.71 | A:Glu | 90 (N) |
| **13** | A:Glu | 84 (O) | 2.84 | A:Gln | 88 (N) |
| **14** | A:Val | 86 (O) | 2.89 | A:Val | 86 (N) |
| **15** | A:Gln | 88 (O) | 3.18 | A:Glu | 84 (N) |
| **16** | A:Gln | 88 (O) | 2.71 | A:Gln | 83 (N) |
| **17** | A:His | 87 (NE2) | 3.31 | A:Glu | 85 (OE2) |
| **18** | A:Glu | 85 (OE2) | 3.31 | A:His | 87 (NE2) |

**Supplementary Table 4.** Confidence metrics for AlphaFold 3 modeled complexes. Interface predicted template modeling (ipTM) and predicted template modeling (pTM) values higher than 0.80 mean that structures represent high-quality predictions. n.a., not applicable.

|  | *ipTM* | *pTM* |
| --- | --- | --- |
| **1x MfpB** | n.a. | 0.81 |
| **1x MfpB – 1x MgGTP** | 0.95 | 0.80 |
| **1x MfpB – 1x GDP** | 0.93 | 0.79 |
| **1x MfpB – 2x MfpD - 1xMgGTP** | 0.89 | 0.90 |
| **1x MfpB – 2x MfpD - 1xGDP** | 0.79 | 0.82 |

**Supplementary Figure 1.**
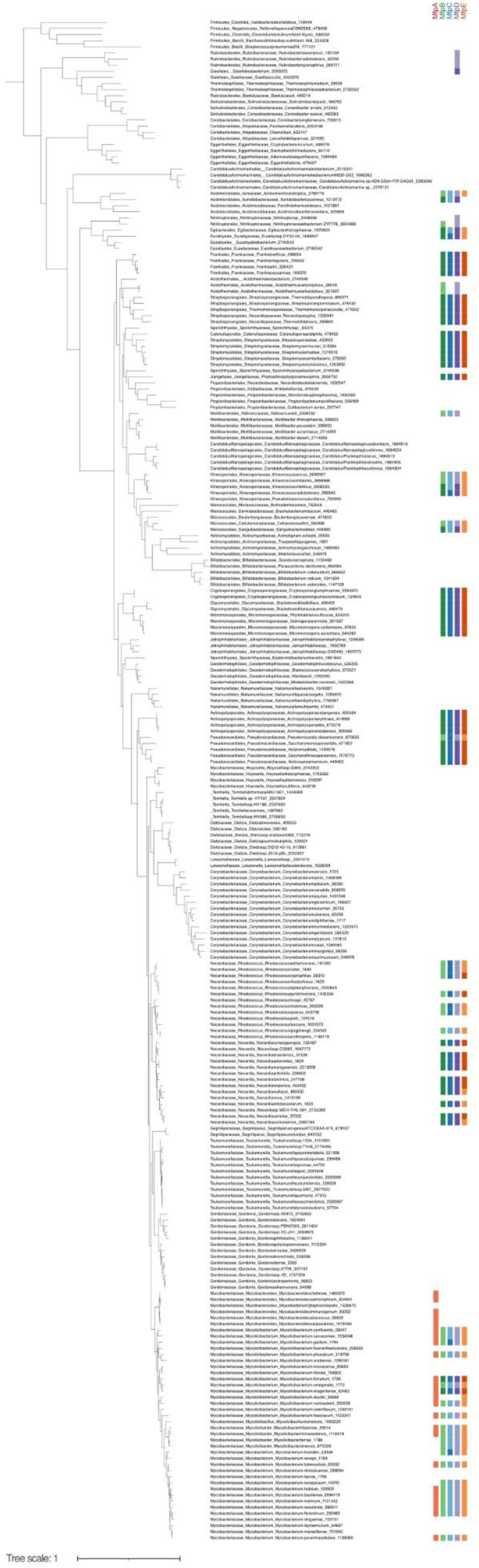
Phyletic pattern of the presence of genes mfpA, mfpB, mfpC, mpfD and mfpE mapped onto a reference phylogeny of the Actinobacteria. A colored square indicates the identification of the gene in at least half of the genomes analysed for the corresponding lineage. A darker shade of the color indicates the identification of more than one copy of the gene per genome. The scale bar represents the average number of substitutions per site. For a summarized version, see Figure 1d.

**Supplementary Figure 2.**
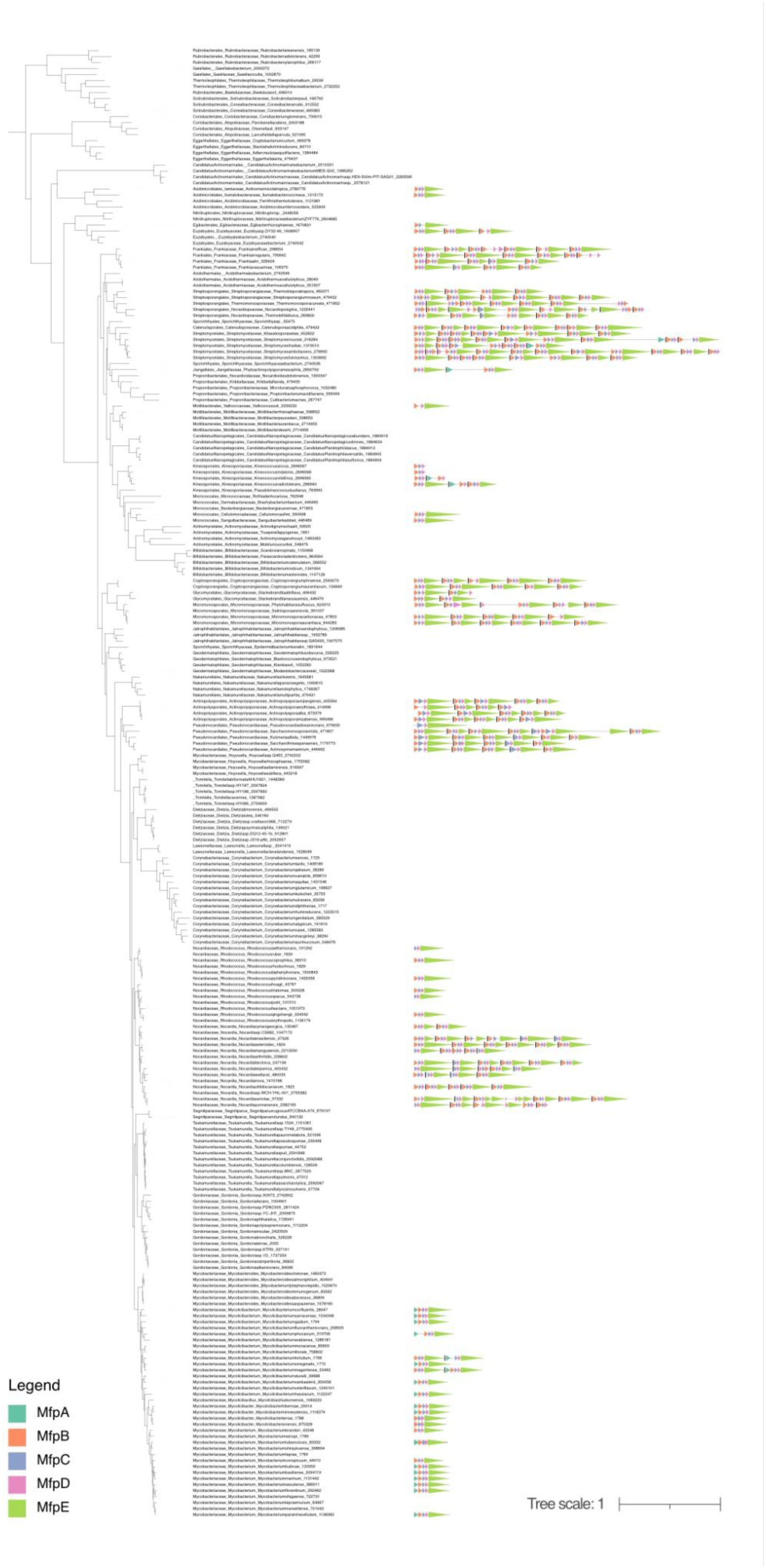
Phyletic pattern of the presence of the mfp conservon mapped onto a reference phylogeny of the Actinobacteria. Multiple conservons identified for the same genome are separated by a vertical black bar. The scale bar represents the average number of substitutions per site.

**Supplementary Figure 3.**
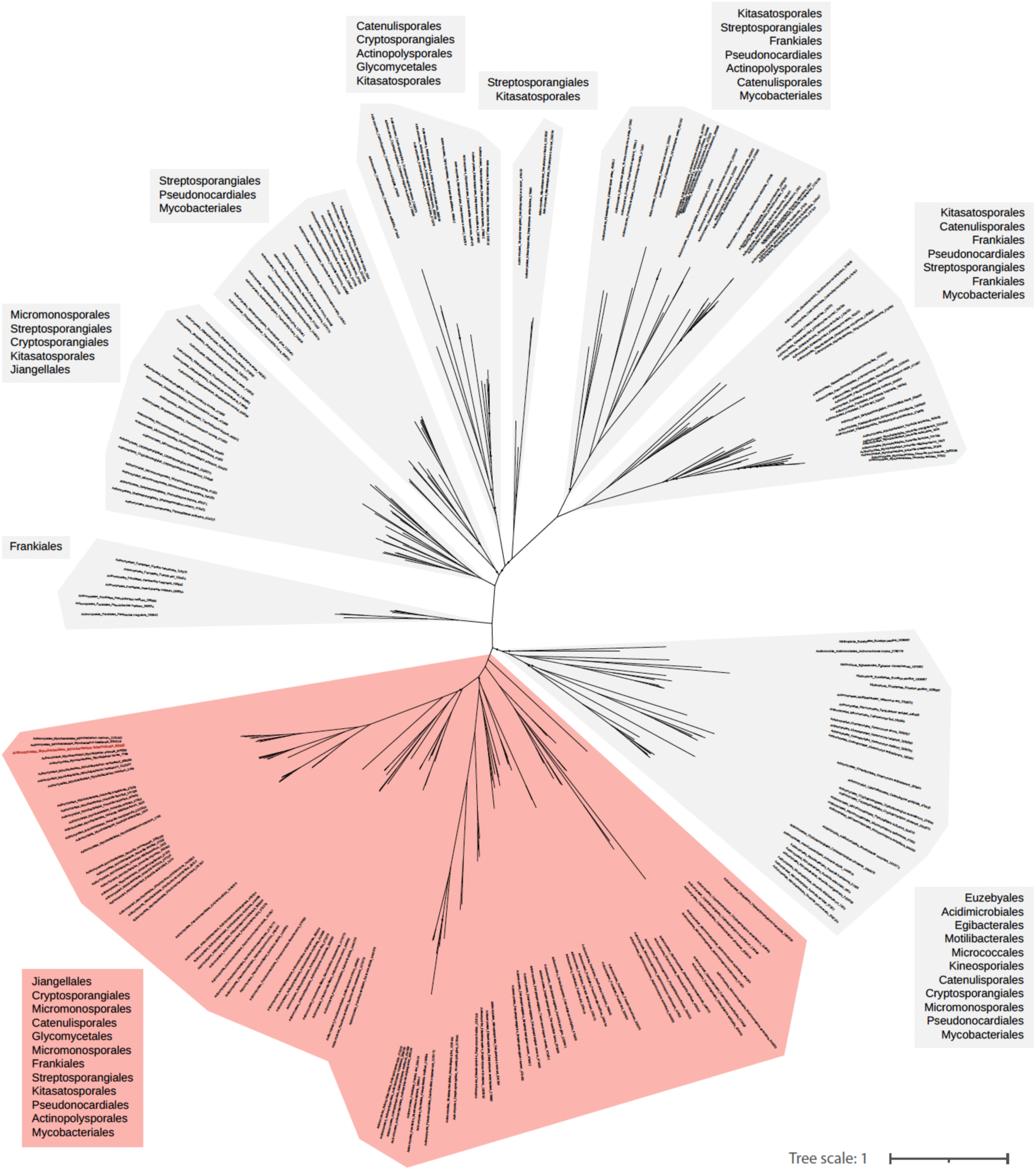
Maximum-likelihood phylogeny of the mfp conservon in Actinobacteria. Monophyletic clades are indicated with separate backgrounds. The largest clade is indicated in pink. Note that the topology of this clade roughly matches that of the reference phylogeny of Actinobacteria. UFB <= 80 are indicated with a dot. The scale bar represents the average number of substitutions per site.

**Supplementary Figure 4.**
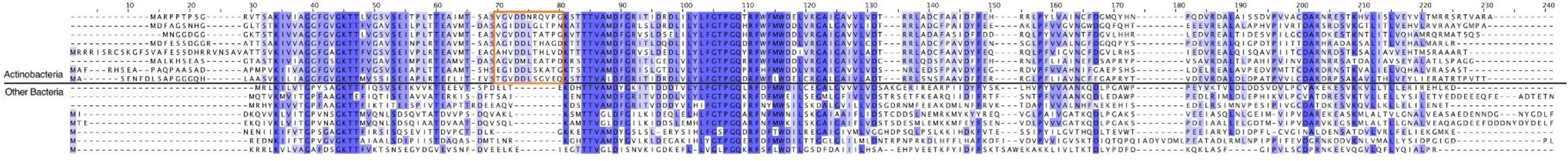
Multiple sequence alignment of MfpB including representative sequences of Actinobacteria and other bacterial phyla. The specific actinobacterial insertion sequence is squared in orange.

**Supplementary Figure 5.**
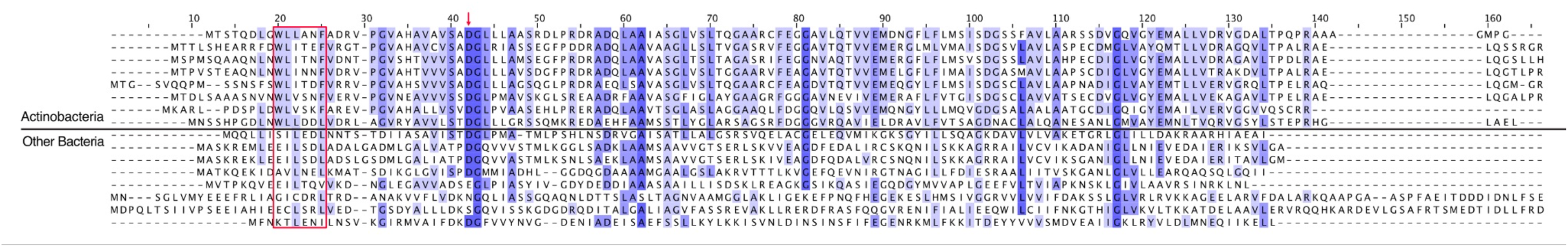
Multiple sequence alignment of MfpD including representative sequences of Actinobacteria and other bacterial phyla. The specific actinobacterial sequence *^12^*WLXXXF^17^ (*Mtb* numbering) sequence is squared in red.

**Supplementary Figure 6.**
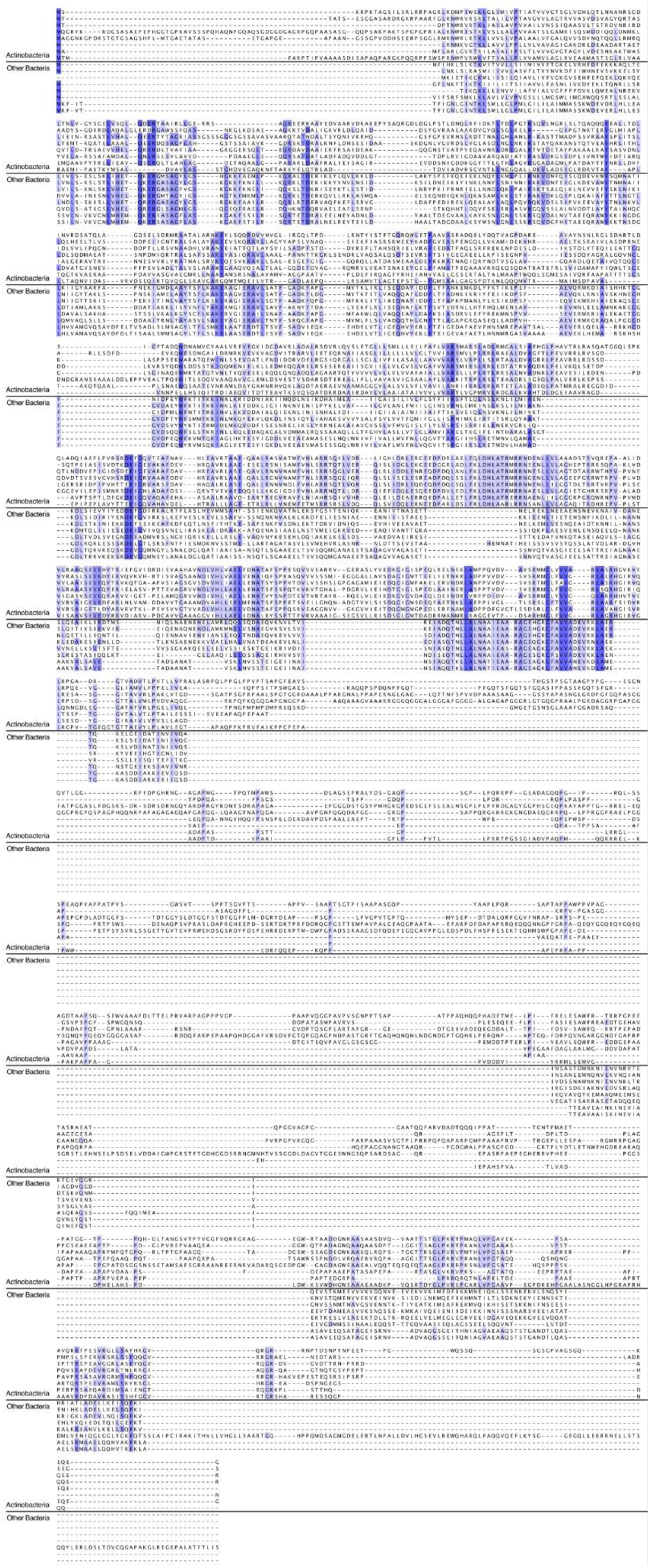
Multiple sequence alignment of MfpE including representative sequences of Actinobacteria and other bacterial phyla.

**Supplementary Figure 7.**
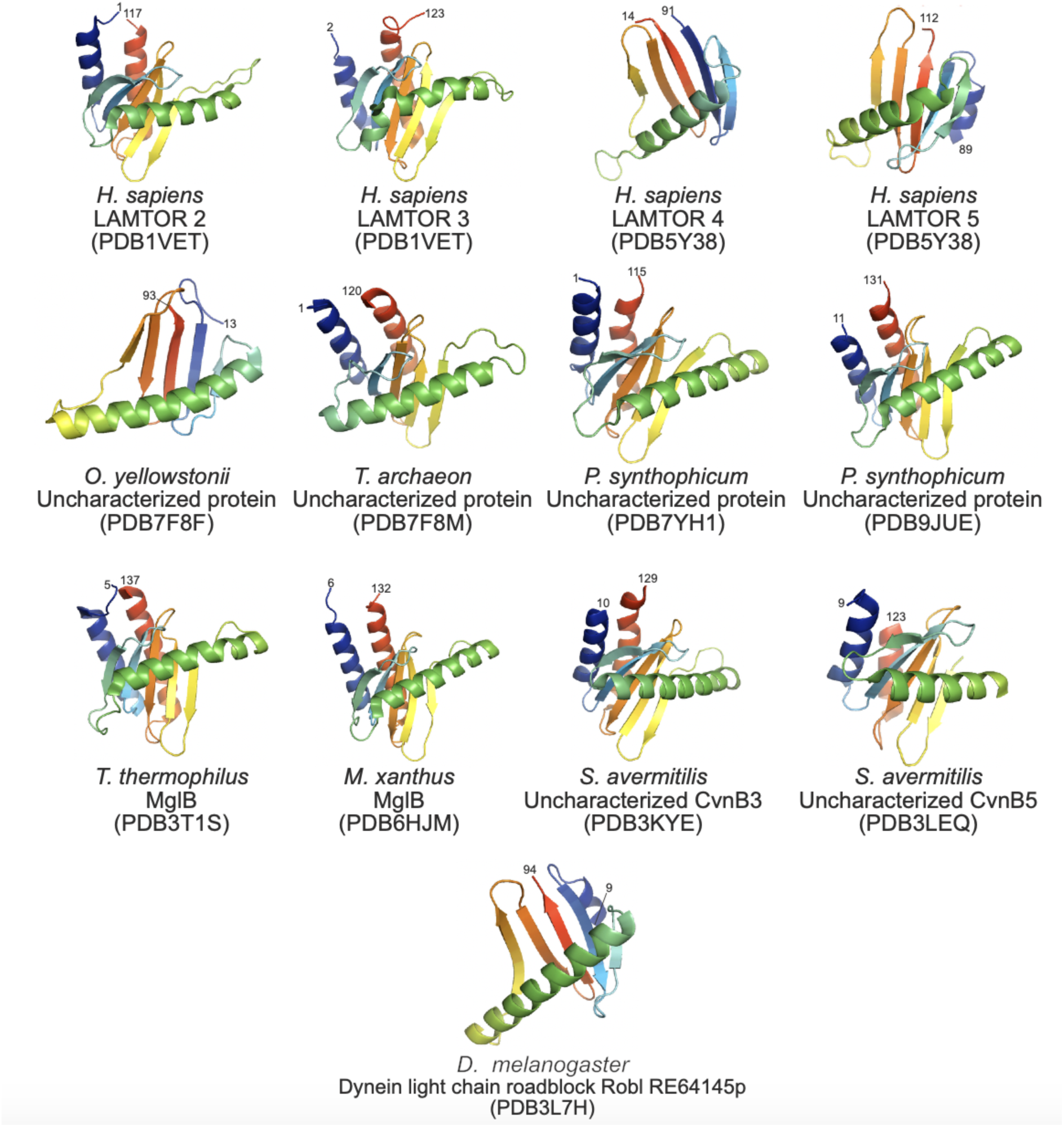
Some of the previously solved crystal structures of Roadblock/LC7 domain-containing proteins. Note that all structures have been reported to be in homo– or hetero-dimeric forms.

**Supplementary Figure 8.**
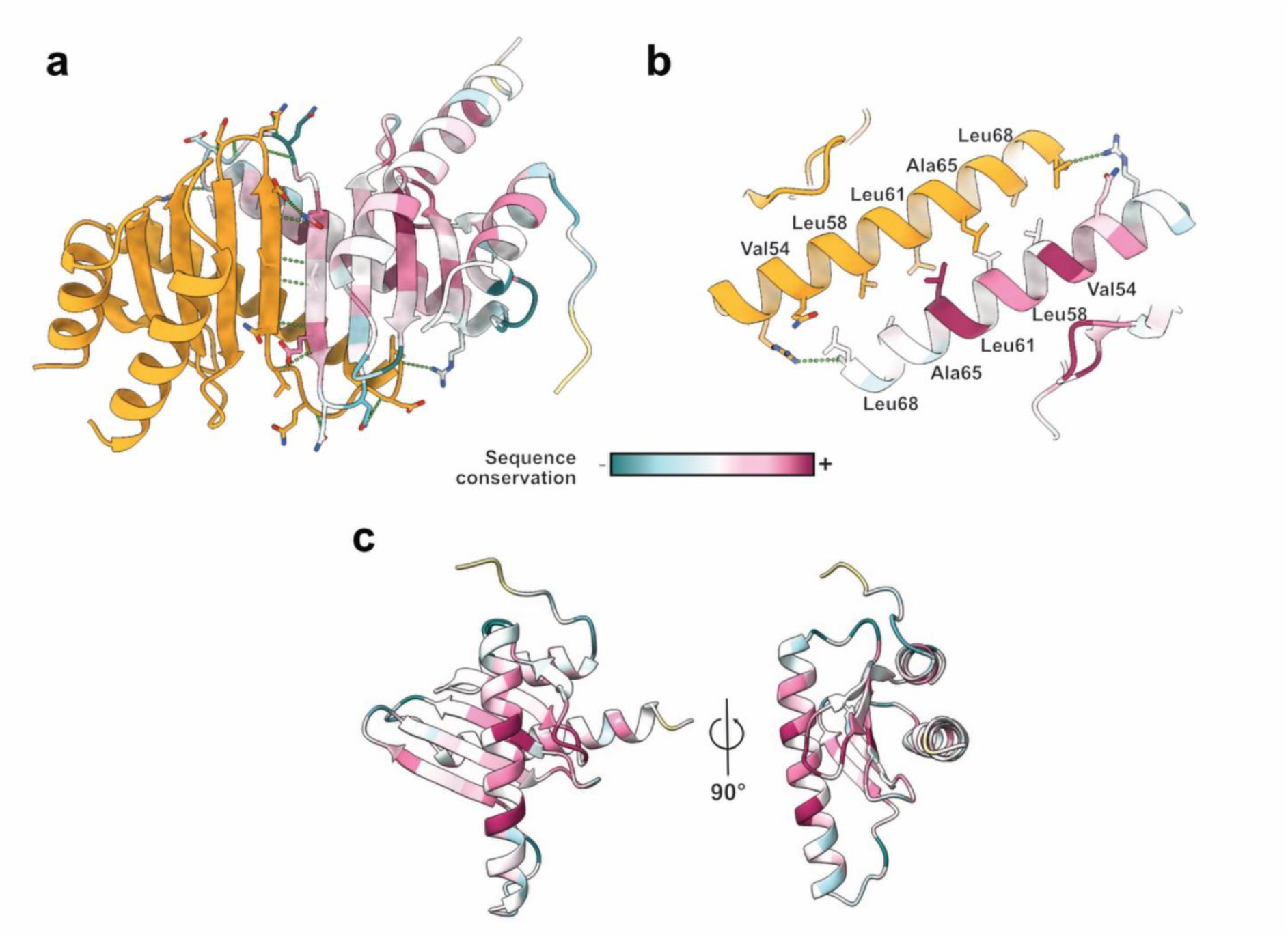
Dimerization interface properties of MfpD. One of the two monomers is colored according to a Consurf analysis to highlight conservation patterns in MfpD-related proteins. Hydrogen bonds are represented by green dashed lines, and all polar residues involved in the polar interactions are displayed in stick format.

**Supplementary Figure 9.**
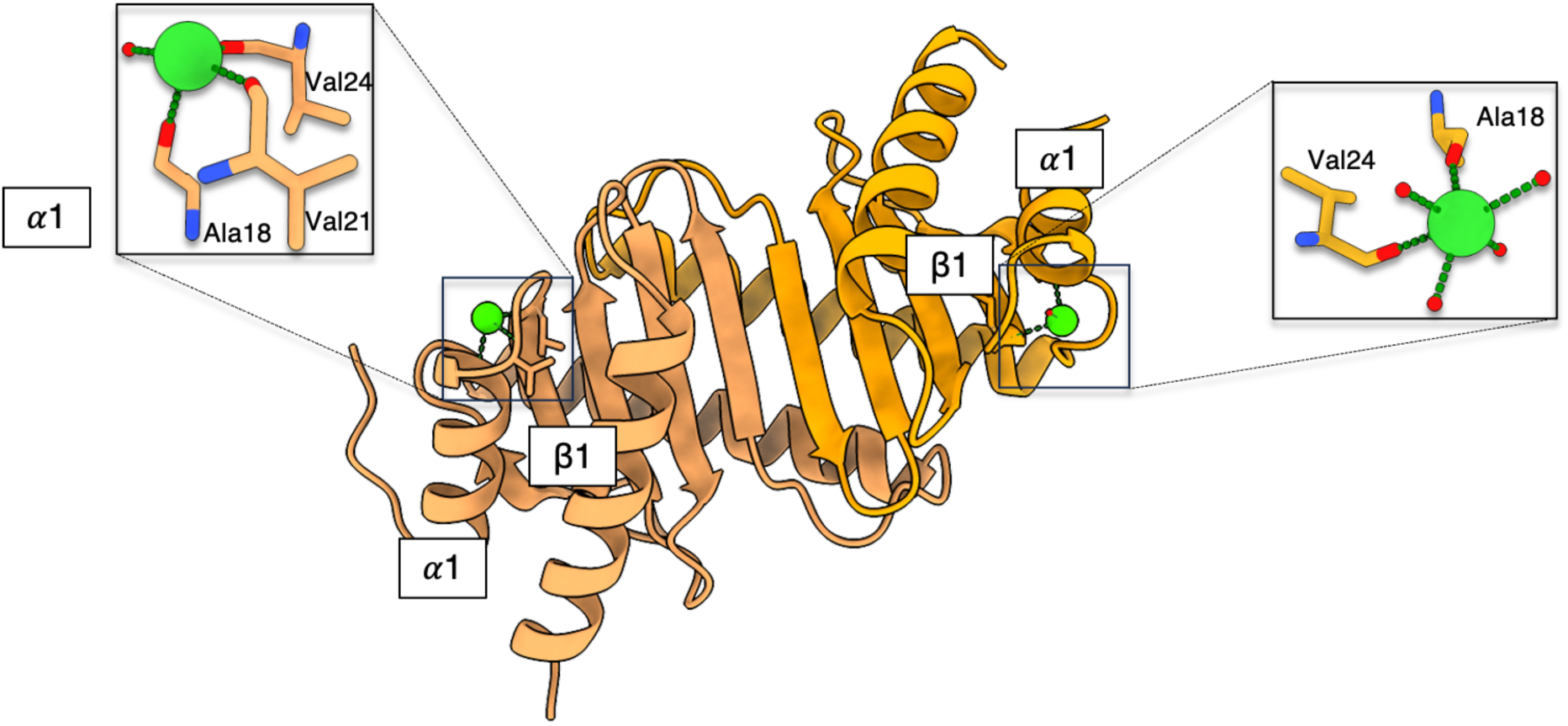
Binding of sodium ions within the MfpD dimer.

**Supplementary Figure 10.**
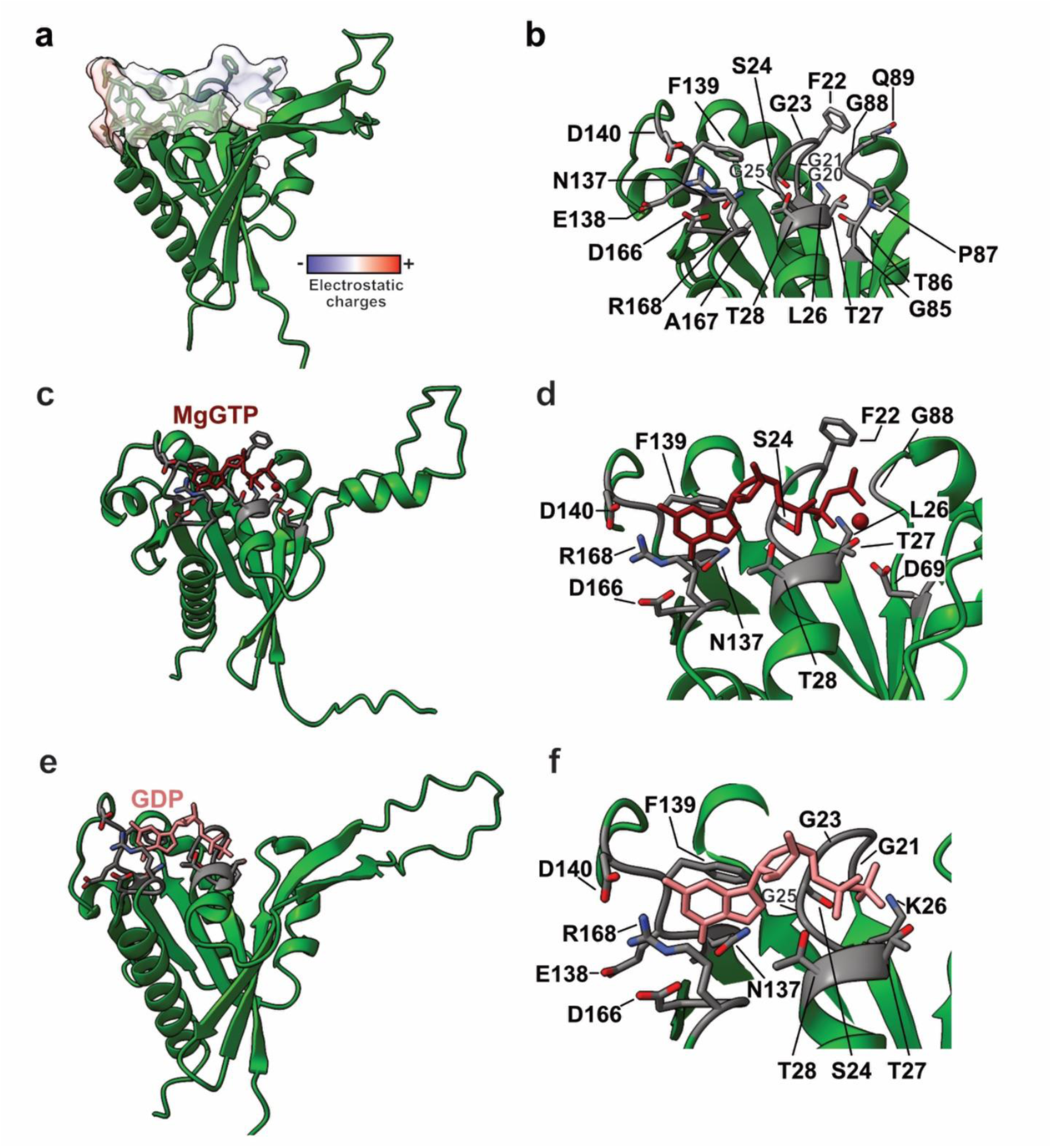
AlphaFold-assisted modeling of Mtb MfpB. AlphaFold3 model of *Mtb* MfpB (***a,b***), *Mtb* MfpB-GTP complex (***c,d***) and *Mtb* MfpB-GDP complex (***e,f***). Zoom on active sites residues in the absence of ligands (***b***) or in the presence of GTP (***d***) or GDP (***f***). The electrostatic charges of the surface of the active site are depicted in ***a***. For simplicity, the unstructured N-terminal sequence is not shown.

**Supplementary Figure 11.**
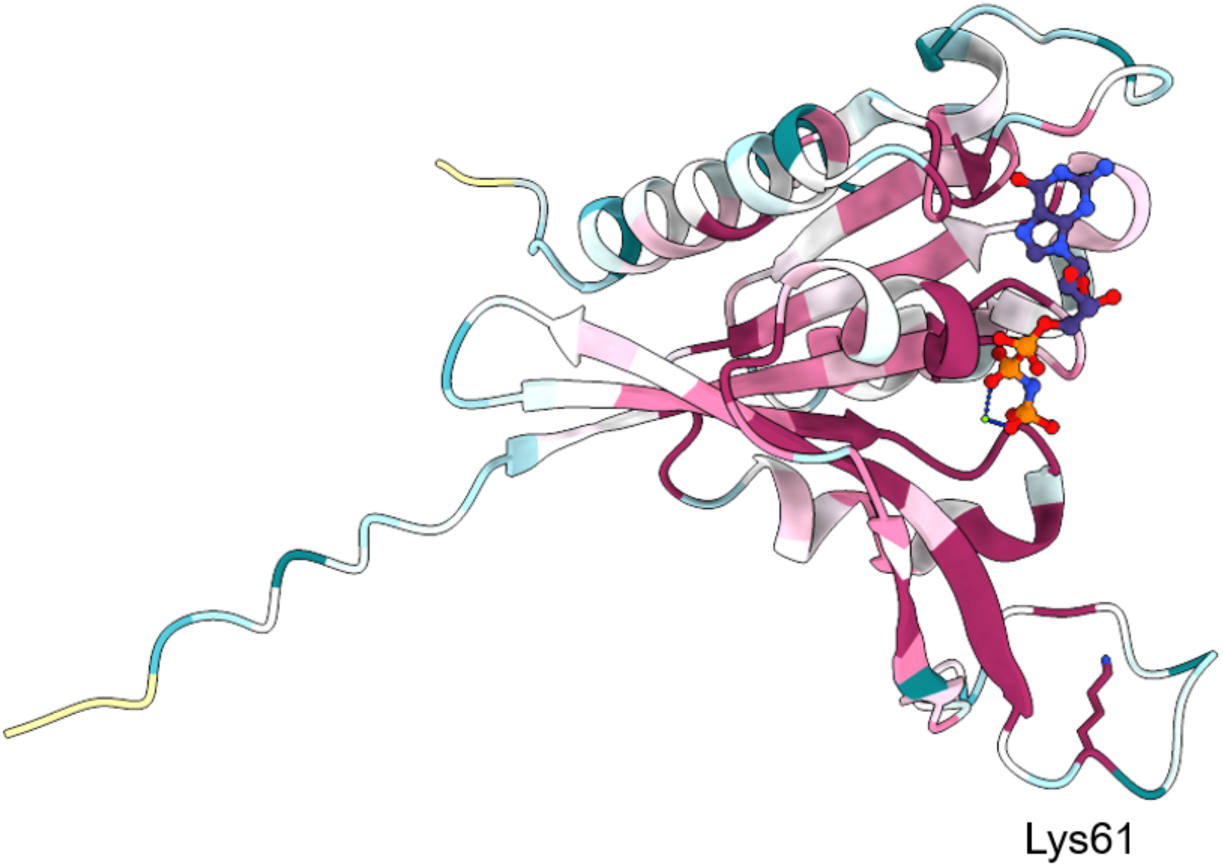
Lysine 61 conservation and position in MfpB model.

**Supplementary Figure 12.**
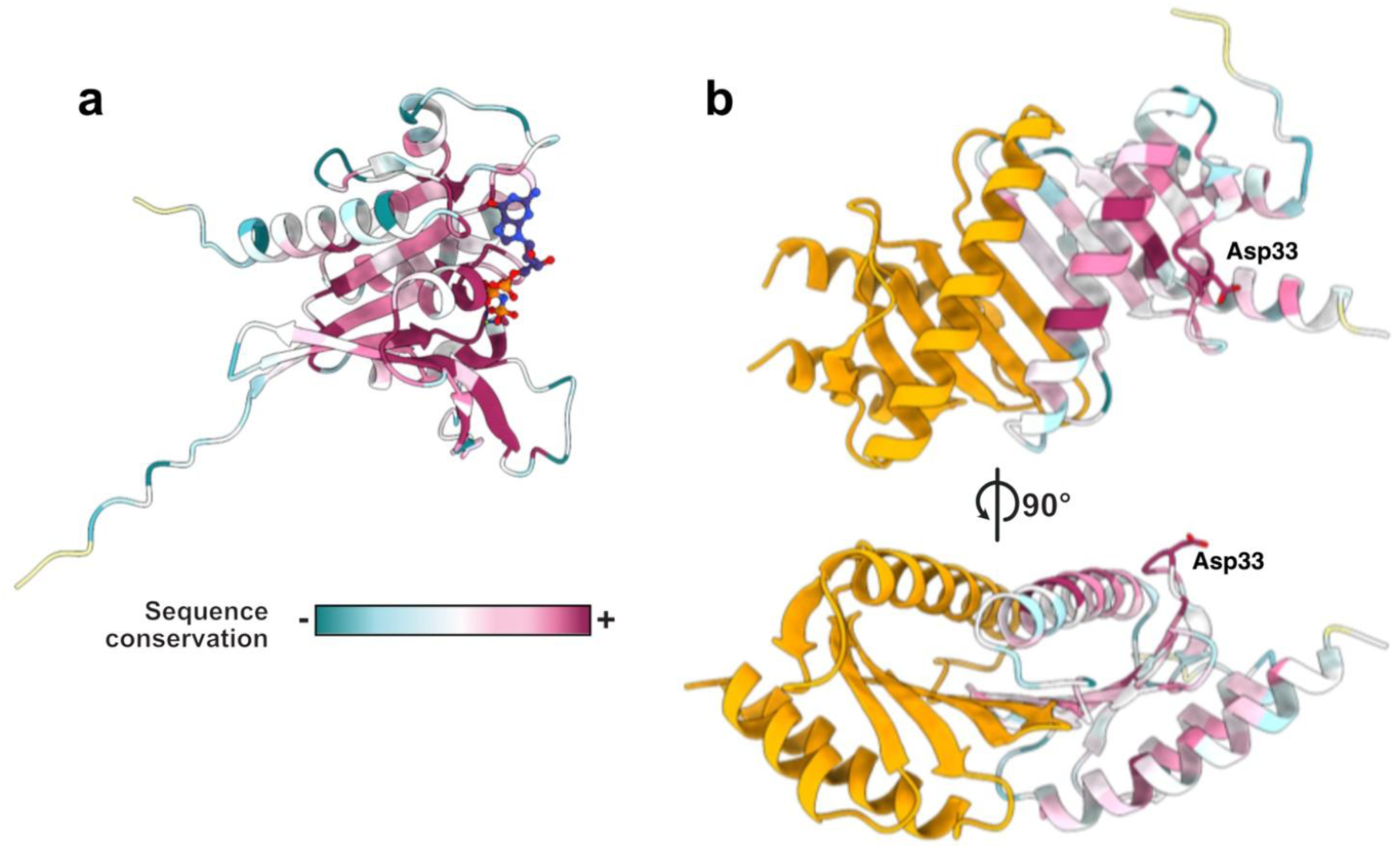
Representation of the conservation level of MfpB (a) and aspartate 33 conservation and position in MfpD dimer (b).

**Supplementary Figure 13.**
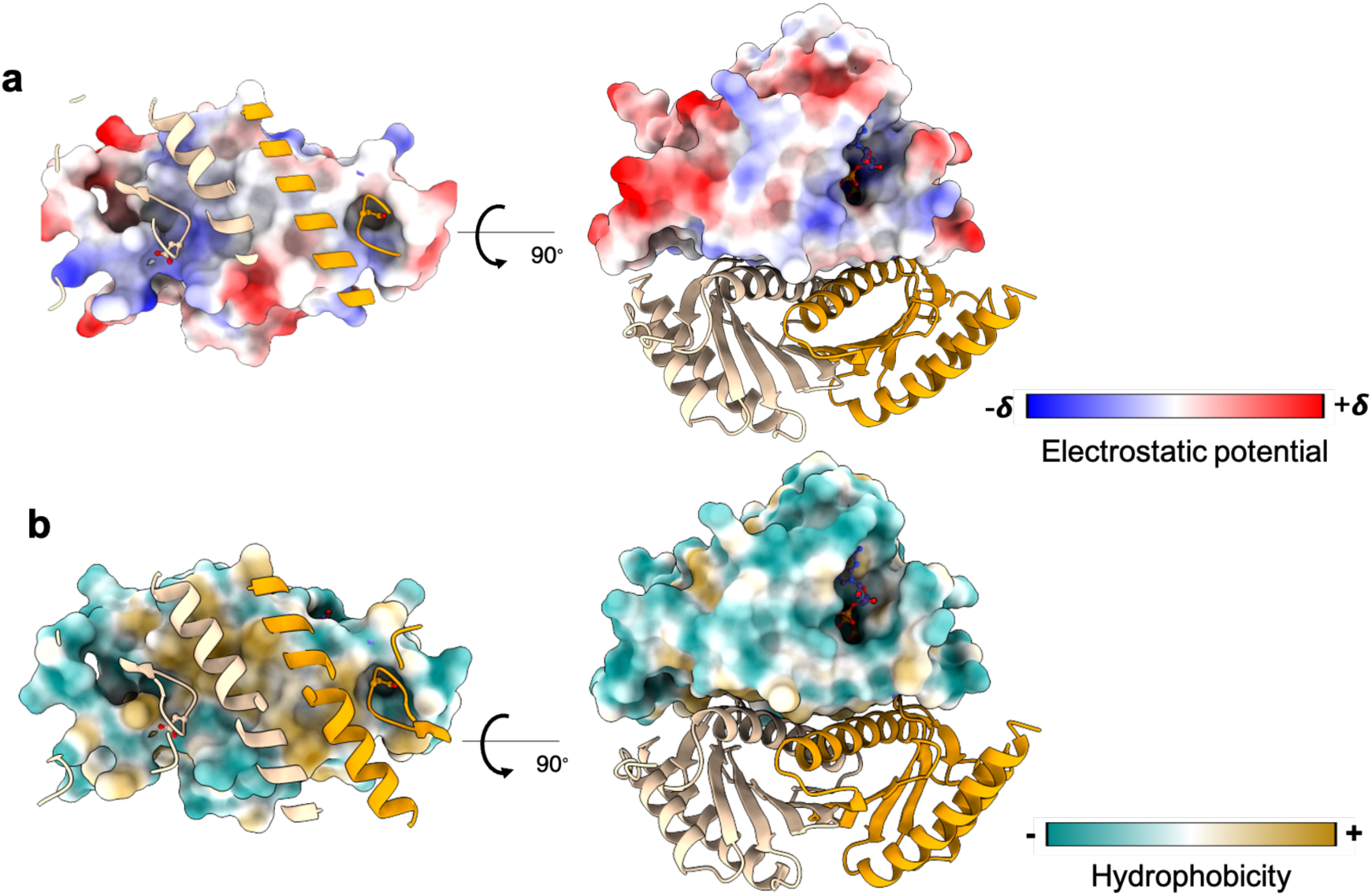
2MfpD-1MfpB complex. ***a.*** Electrostatic potential surface and ***b.*** hydrophobicity potential surface for MfpB.

**Supplementary Figure 14.**
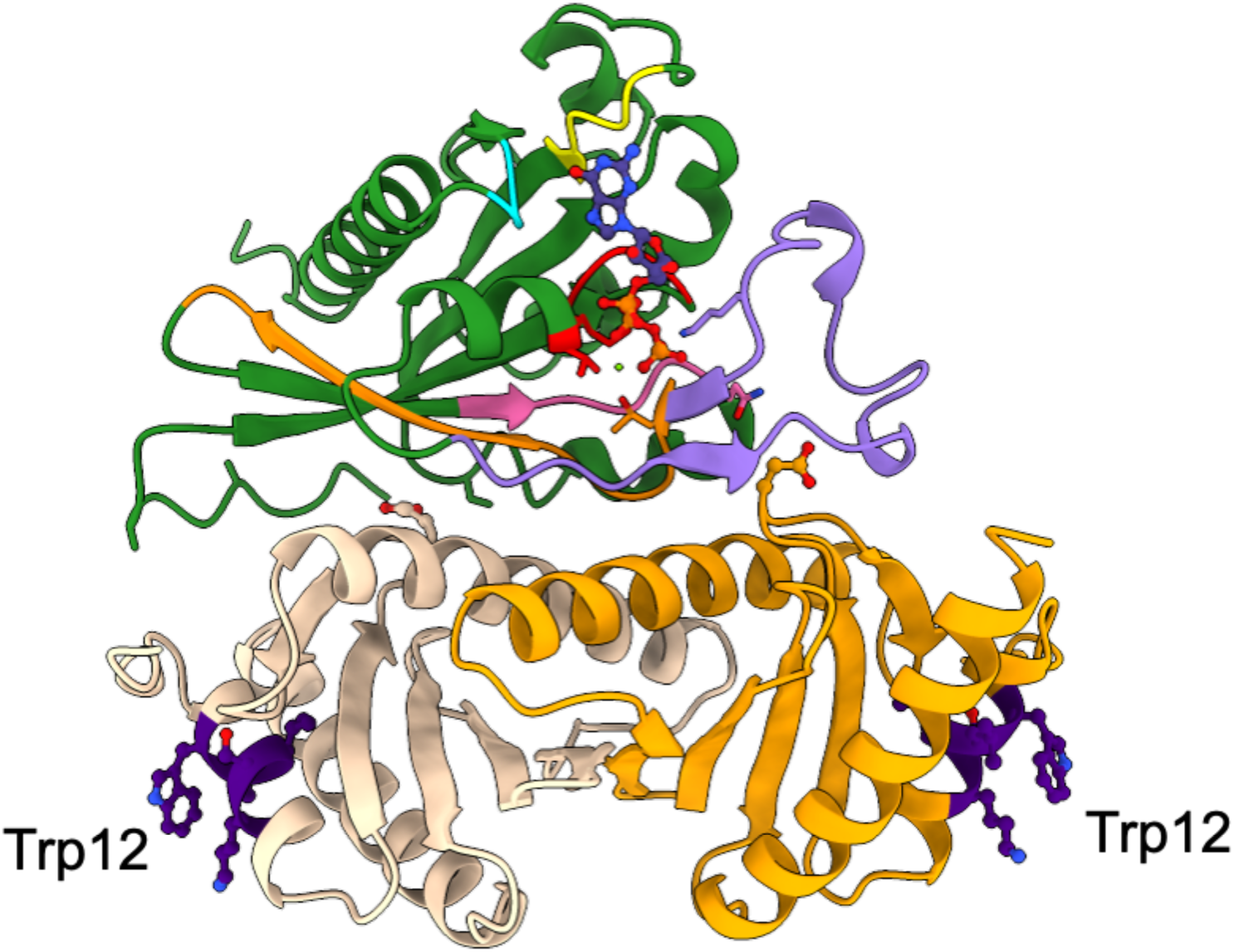
Positioning of *^12^*WLVSKF^17^ sequence (in dark purple) in MfpD.

**Supplementary Figure 15.**
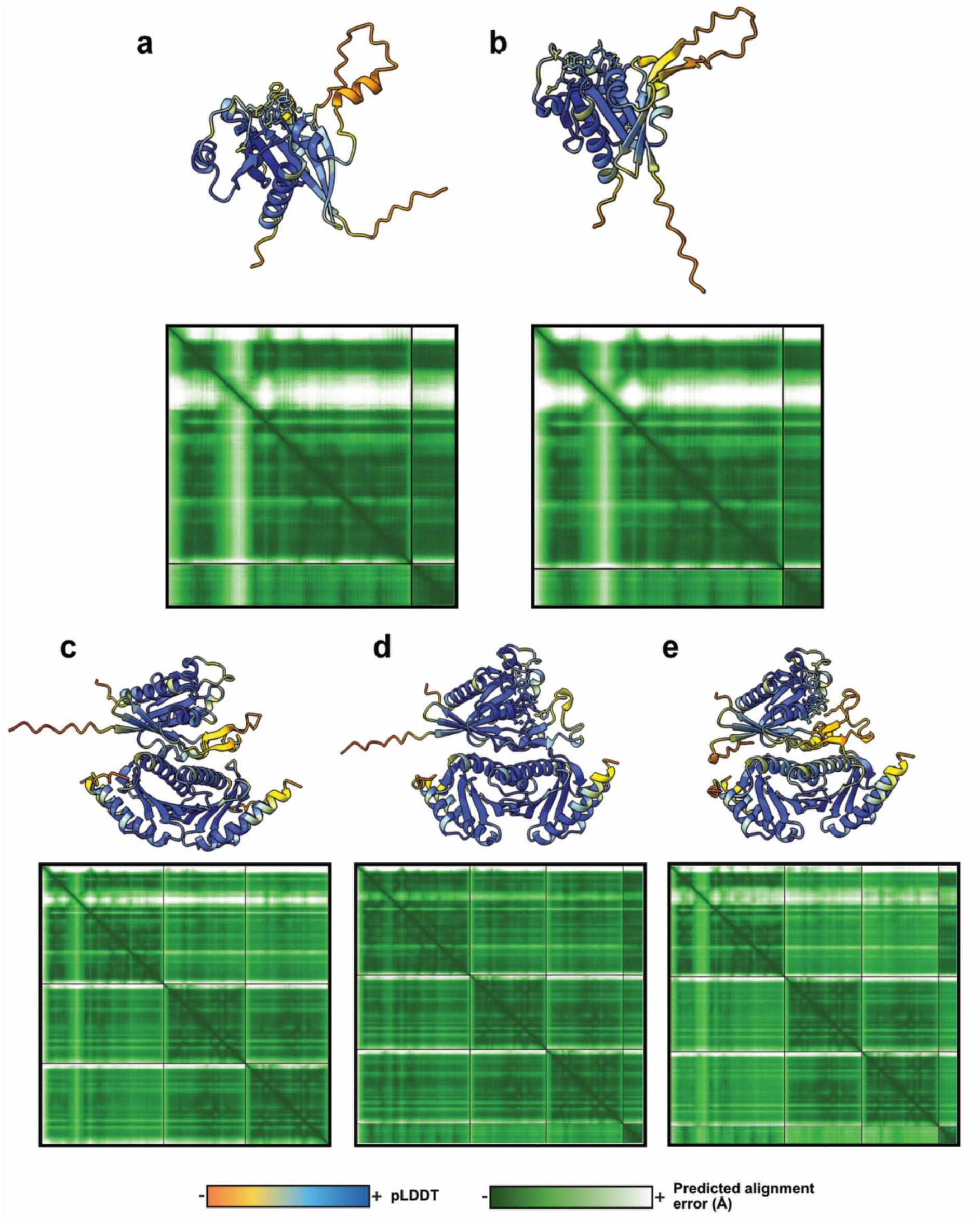
AlphaFold3 prediction confidence levels. Three-dimensional structures of MfpB-MgGTP (a), MfpB-GDP (b), MfpD-MfpB (c), MfpD-MfpB-MgGTP and (d) MfpD-MfpB-GDP models colored per-residue confidence level and corresponding residue-residue alignment confidence plots.

## Notes

### Competing Interest Statement

The authors have declared no competing interest.

### Summary of Updates

Some titles, one miss Sodium ion and the affiliation of my colleagues

